# Skeletal Muscle Specific PolG Dysfunction Activates the Integrated Stress Response and Promotes a Cachexia like Phenotype

**DOI:** 10.1101/2024.06.02.597064

**Authors:** Simon T Bond, Emily J King, Shannen M Walker, Christine Yang, Yingying Liu, Kevin H Liu, Aowen Zhuang, Aaron W Jurrjens, Haoyun A Fang, Luke E Formosa, Artika P Nath, Sergio Ruiz Carmona, Michael Inouye, Kevin Huynh, Peter J Meikle, Nadeem Elahee Doomun, David P de Souza, Danielle L Rudler, Anna C Calkin, Aleksandra Filipovska, David W Greening, Darren C Henstridge, Brian G Drew

## Abstract

During mitochondrial damage, information is relayed between the mitochondria and nucleus to coordinate precise responses to preserve cellular health. One such pathway is the mitochondrial integrated stress response (mtISR), which is specifically activated by mitochondrial DNA (mtDNA) damage. However, the causal molecular signals responsible for this activation remain elusive. A gene often associated with mtDNA mutations/deletions is *Polg1*, which encodes the mitochondrial DNA Polymerase γ (PolG). Here, we describe what is to our knowledge the first conditional muscle specific model of PolG mutation, and demonstrate a rapid development of mitochondrial dysfunction and cachexia in these animals from ∼5 months of age. Detailed molecular profiling of muscles demonstrated robust activation of the mtISR, likely mediated upstream by the HRI/DELE1 stress pathway. This was accompanied by striking alterations to the mitochondrial folate cycle and its metabolites, strongly implying imbalanced folate intermediates as a previously unrecognised pathology driving the mtISR in mitochondrial disease.

## INTRODUCTION

Mitochondria are small, double membraned organelles found in most eukaryotic cells, often in their thousands. Their cellular roles include the regulation of several processes including apoptosis, redox balance, inflammation and ATP production. Mitochondria are unique in that they harbour their own genome (mtDNA), which is small and circular, and encodes for 37 of the approximately 1500 genes required for efficient mitochondrial function ^1–3^. Of the 37 genes, 22 are transfer RNAs (tRNA), 2 are mitochondrial ribosomal RNAs (rRNA), and 13 are protein coding genes critical for optimised functioning of the mitochondrial electron transport chain (ETC) ^4–7^. The remainder of the ∼1500 required mitochondrial proteins are transcribed by the nuclear genome, translated in the cytosol, then imported into the mitochondria. Each mitochondrion can harbour several copies of mtDNA, where it is replicated and maintained by dedicated transcriptional machinery including Mitochondrial transcription factor A (TFAM), DNA polymerase gamma (PolG), helicase Twinkle (TWNK), Mitochondrial single-strand binding protein (SSBP1) and Mitochondrial RNA polymerase (POLRMT)^7^. PolG is proposed to be the sole DNA polymerase found in the mitochondria and is essential for the replication and maintenance of mtDNA^8^. The mature PolG enzyme consists of three components: the catalytic PolG1 protein, and two units of the accessory subunit PolG2. The PolG enzyme has three main enzymatic actions; DNA polymerase activity, DNA exonuclease activity (proofreading), and base excision repair^9^.

Mutations to mtDNA are often a cause of mitochondrial dysfunction and subsequent mitochondrial disease (MD) ^10^. Indeed, the most common genetic cause leading to an increase in mtDNA mutations, is coding mutants in the *Polg1* gene ^11^. *Polg1* mutations can, amongst other defects, lead to a dysfunctional PolG enzyme complex that lacks the ability to efficiently replicate or proofread mtDNA, ultimately leading to an accumulation of mtDNA mutations/deletions, mtDNA depletion and subsequent mitochondrial dysfunction ^11–13^. PolG induced diseases vary dramatically in their presentation, and often develop unpredictably from birth through to later life. PolG pathologies are various in their presentation and lead to degenerative muscle, neurological and metabolic diseases ^12–16^. PolG induced mtDNA mutations/deletions mostly occur randomly across the mitochondrial genome, although the major arc (between the two origins of replication) attracts a major part of this mutational burden ^17,18^. Often, the location of the mutation is unrelated to the development of clinical symptoms, where instead it is the burden of overall mutational load in a given tissue/cell that is the pathological switch ^19,20^. Cells that harbour both WT and mutant variants of mtDNA are described as being heteroplasmic, where the cell can maintain mostly normal function so long as the ratio of mutant versus WT copies remains below a given threshold ^21^. However, once the ratio exceeds that threshold, disease pathologies rapidly develop ^19–21^.

Much of our prior understanding of the mechanisms underlying PolG mediated disease, has been generated from human studies in individuals carrying SNPs in the *Polg* coding sequence, and from mouse models that harbour a whole body insufficiency in PolG exonuclease activity, known as the mitochondrial mutator mouse - of which two independent models currently exist ^12,13^. These transgenic mice were generated by base substitution of an aspartate to alanine residue at site 257 (D257A) of the PolG peptide, which falls within the conserved exonuclease domain. The D257A Mutator mice subsequently lack PolG exonuclease activity and thus accumulate mtDNA mutations over time ^12,13^. Homozygous D257A PolG mice present with multiple disorders by early adulthood including the progressive loss of body mass, skeletal muscle and bone mass, amongst other ailments such as cardiomyopathy, premature aging, kyphosis, impaired mobility and ultimately premature death ^11–14,22^. As expected, the organs most affected in these mice are those that are heavily reliant on mitochondrial energy production such as the brain, muscle and heart. However, the temporal and tissue-specific aspects of disease progression in the mutator mouse has been difficult to dissect due to its developmental nature. For example, with the described heart failure that develops in these mice ^23^, it is difficult to determine how much of this pathology is due to intrinsic mitochondrial defects in cardiomyocytes, compared to systemic issues relating to metabolic dysregulation, neurological degeneration and skeletal muscle atrophy. Given this predicament, it is important to advance our understanding of how increasing mtDNA mutation load affects these tissues individually, and how this subsequently drives complications.

Herein, we describe what is to our knowledge the first conditional mouse model that allows for specific removal of the PolG exonuclease domain, resulting in tissues that express a truncated PolG protein with all elements of function intact – other than the proof-reading/repair capacity. This mouse can be crossed with cre-recombinase expressing mice to generate models with tissue-specific manipulation of PolG activity. We validated this model in an inducible, post-developmental skeletal muscle specific model (ACTA1-cre-ERT2), which presented with a systemic phenotype from approximately 5 months post-recombination. Importantly, not all features of the whole body PolG mutator mouse are recapitulated in our model, suggesting that many of the features observed in the global mutator mouse either arise from developmental defects, differential aetiology, or from tissue crosstalk arising from mitochondrial dysfunction in other tissues. Importantly, we also use this mouse to reveal fundamental insights into our understanding of the mitochondrial integrated stress response (mtISR), where evidence presented herein suggests that mtDNA deletions induced by PolG dysfunction, lead to alterations in the mitochondrial folate cycle which likely contribute to the chronic activation of the mtISR pathway.

## RESULTS

### Generation and Validation of a Muscle Specific PolG Exonuclease Mutant

In order to understand tissue-specific PolG phenotypes in the absence of confounding deficits from other organs, we sought to develop a Cre-Lox mediated model in C57BL/6J mice, which allowed for temporal and tissue specific manipulation of PolG exonuclease (DNA proofreading) activity. We aimed to specifically interrupt PolG exonuclease activity to induce mtDNA damage (deletions/mutations), which is often the prevailing insult leading to disease phenotypes in a high proportion of humans who have pathogenic polymorphisms in the PolG gene.

To generate this unique mutant, it was necessary to remove all or some of the exonuclease coding region of the PolG mRNA (encoded by exons 2-6 in the mouse), whilst allowing for the remaining mRNA components to be preserved in their expression and translation. In silico analysis of the mRNA sequence revealed that targeting of exons 4 and 5 of the PolG transcript would truncate ∼50% of the exonuclease domain, but should allow for the remainder of the mRNA to remain in frame. Hence, we flanked exon 4 and 5 with LoxP sites using CRISPR and cross-bred mutant mice with wildtype C57BL/6J mice for 5 generations, before crossing to Cre-recombinase expressing mice. In this study we chose to study the impact of PolG exonuclease deletion in skeletal muscle using the ACTA1-cre-ERT2 mouse (**Figure 1A**). This model allows for temporal and skeletal muscle specific deletion via expression of Cre-recombinase from the ACTA1 promoter, and Tamoxifen inducible Cre-activation via the ERT2 domain. We focussed on skeletal muscle PolG dysfunction due to the previously described degenerative phenotype (ragged red fibres) observed in global mutator (PolG D257A transgenic) mice^16^, and the prominent role that skeletal muscle plays in mitochondrial disease phenotypes. Mice were bred to generate two groups of mice (fl/fl and fl/fl-Cre) that were further divided into groups that either received Tamoxifen (80mg/kg) in oil, or oil alone resulting in 4 groups in total; fl/fl-OIL, fl/fl-TAM (PolG^cont^), fl/fl-cre-OIL and fl/fl-Cre-TAM (PolG^mut^). It is important to highlight that these mice were allowed to develop normally until 8 weeks of age, after which tamoxifen was administered to generate mutants in adolescent mice – resulting in a post-developmental PolG mutant model. Cohorts were then studied for two different time periods: 4 months (referred to from this point as “short-term”) and 12 months (referred to from this point as “long-term”) (**Supplemental Figure 1A**).

**Figure 1:**
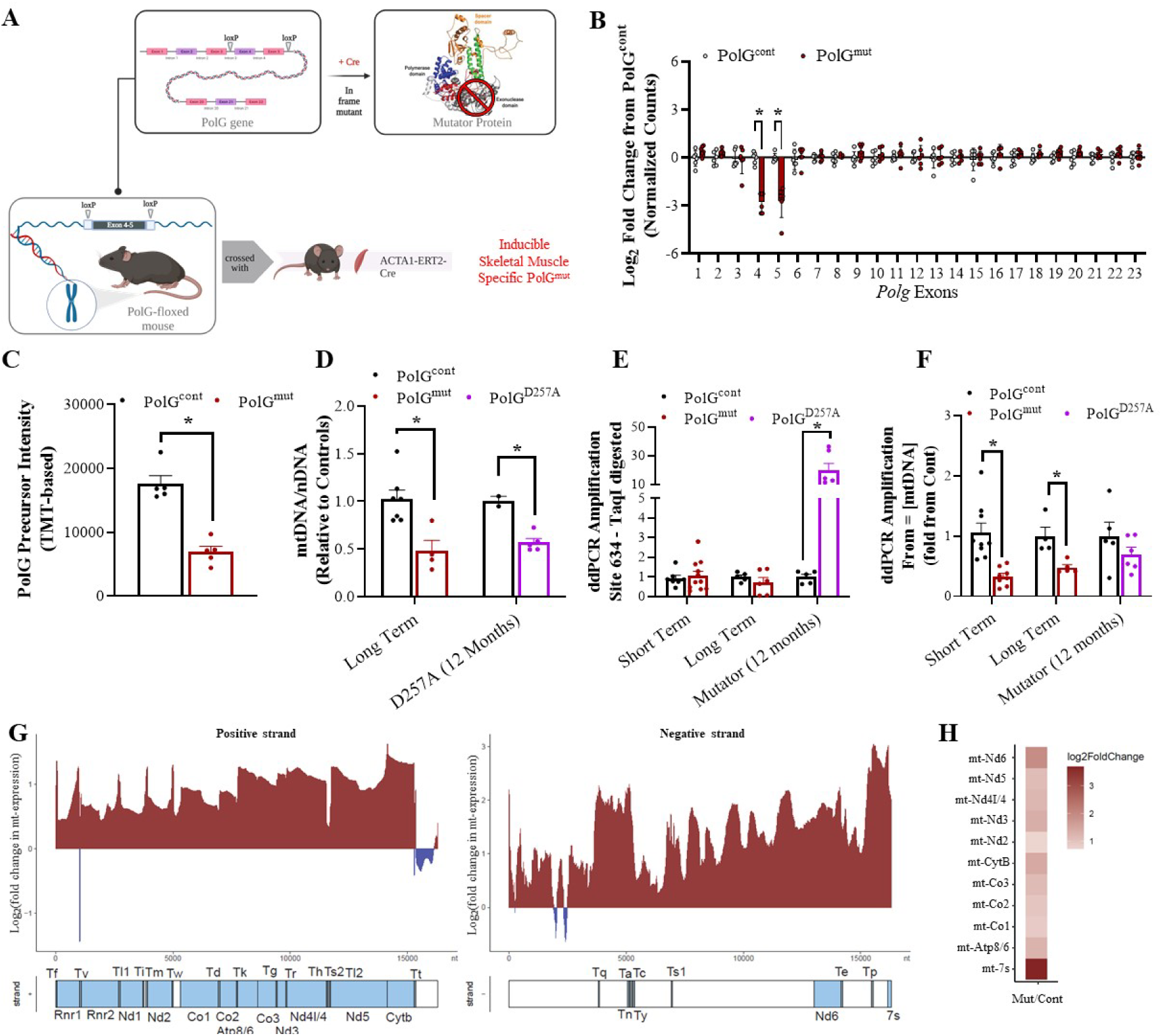
Generation of a Post-developmental Muscle Specific PolG Exonuclease Mutant Mouse. **(A)** Schematic outlining the design of the floxed PolG mouse and the genetic regions (exons 4&5) that are targeted for deletion by cre-recombinase. Crossing the floxed mouse with the ACTA1-cre-ERT2 mouse generates a skeletal muscle specific loss of these exons resulting in an in-frame mutant lacking critical regions of the exonuclease domain. **(B)** Change in abundance of each exon across the *Polg1* gene in skeletal muscle, as determined by RNA-sequencing, presented as Log_2_ fold change in exon abundance in PolG^mut^ (red bars) compared to control (PolG^cont^, black bars) mice. **(C)** PolG precursor ion intensity (protein abundance) from proteome analysis of skeletal muscle of PolG^cont^ and PolG^mut^ mice (n=5). **(D)** Ratio of mitochondrial DNA (mtDNA) to nuclear DNA (nDNA) (normalised to control) in skeletal muscle of PolG^cont^ and PolG^mut^ mice at 12 months of age (12 months) and in the skeletal muscle of the global D257A PolG mutator mouse at 12 months of age (aD257A – 12 months). **(E)** Mutation rate (fold change from control) as determined by RMCA and ddPCR in skeletal muscle mtDNA of PolG^mut^ and D257A Mutator mice in short or long-term cohort compared to their respective controls. **(F)** Amplification rate (fold change from control) of a “control” mtDNA region as determined ddPCR from equivalent amounts of skeletal muscle mtDNA in PolG^mut^ and D257A Mutator in short or long-term cohorts compared to their respective controls. (**G**) Log_2_ fold change of mapped RNA sequence reads across the mtDNA positive and negative strands and (**H**) heatmap representing Log_2_ fold change of mitochondrial RNA transcript abundance in PolG^mut^ compared with PolG^cont^, as determined by RNA-sequencing in skeletal muscle. Data are presented as mean ± SEM, n=5-10, * denotes p-value <0.05 between control and mutant samples as determined by unpaired, two-tailed Mann-Whitney t-tests.

Given that this was the first model generated using this conditional PolG exonuclease mutant mouse, we sought to test key aspects to confirm its validity. We first set out to demonstrate that skeletal muscle specific deletion of exon 4 and 5 was successful, and that this did not result in complete loss of PolG mRNA due to non-sense meditated decay. Using RNA-sequencing data we demonstrate robust reductions (Log_2_ fold = -4; ∼12-16 fold) in counts across both exons 4 and 5 of the PolG mRNA transcript in PolG^mut^ mice, with no evidence of reduction in any other exon (**Figure 1B).** Using qPCR, we show that a significant reduction in exon 4-5 abundance was only observed in the skeletal muscle of PolG^mut^ mice, and not in other tissues including heart and white adipose tissue (**Supplemental Figure 1B**). Together, these data provide evidence that the PolG mRNA transcript in mutant mice is deleted for exons 4 and 5 in skeletal muscle, but retains all other components.

Having demonstrated that the *PolG* mRNA was depleted of exons 4-5 in our model, but otherwise intact, we next wanted to establish that the truncated *PolG* mRNA transcript could indeed generate a protein product, and that we hadn’t inadvertently generated a knockout (KO) of PolG protein. To do this we performed TMT-based and label-free mass spectrometry proteomics to determine PolG abundance in control and mutant skeletal muscle. This approach initially proved challenging due to the limited number of potential tryptic sites within the PolG sequence. However, when using a mitochondrial enrichment approach in combination with TMT-labelling and peptide fractionation strategy with high sensitivity proteomics, we were able to confidently analyse unique/razor peptide from the PolG protein from n=5 animals/group (**Supplemental Figure 1C&D**). Here, we detected PolG in PolG^mut^ muscle (n=5), which demonstrated an approximately 50% (p<0.0079) decrease in PolG (relative to PolG^cont^ mice, n=5) expression based on identified precursor ion intensity (**Figure 1C**). These data collectively demonstrate that our mouse model generates a protein product from the truncated mRNA, and does not represent a KO model of PolG.

With the confidence that our model was generating the appropriate truncated mRNA and detectable levels of a PolG protein, we sought to confirm whether this mutant PolG protein had retained its ability to interact with putative binding partners and perform necessary functions including mtDNA replication. With regards to binding partners, often if one key factor of the mitochondrial transcriptional complex is deleted, other co-factors are also significantly reduced in their abundance – resulting in loss of mtDNA replication activity ^24,25^. Given this finding, we analysed proteins from PolG^mut^ muscle proteome for their abundance of known PolG binding partners (SSBP, POLRMT, TFAM, TWNK) to infer that the mtDNA replication machinery was indeed intact. These data demonstrate that there was some alteration in the abundance of each of these binding partners, with SSBP1, POLRMT and TWNK all being slightly elevated in PolG^mut^ muscle, whilst TFAM was notably reduced (**Supplemental Figure 1E & 1F**). Accordingly, these data suggest that although there were some compensatory effects in our mutant PolG model, the mtDNA replication machinery appeared to be otherwise replete. To investigate this further, we quantified the abundance of mtDNA in whole muscle from PolG^mut^ and PolG^cont^ mice at 12 months of age, alongside heterozygous 12 month old mice from the previously described global D257A PolG mutator mouse (which does not have a major replication deficit)^12^. These data confirm that both models demonstrated the same partly reduced mtDNA abundance compared to control mice (**Figure 1D**), implying that our skeletal muscle specific PolG^mut^ model behaved similarly with regards to mtDNA replication as did the well described global PolG mutator model.

Another important validation of this model, is determining whether removal of the exonuclease domain led to alterations in the integrity of mtDNA (mutations/deletions) in skeletal muscle. To investigate this we performed the Random Mutation Capture (RMC) assay^26^, coupled with digital droplet PCR (ddPCR) which together can be used to estimate mutation and deletion abundance^26^ in the mtDNA. As a positive control for this assay, we used mtDNA isolated from muscle of the global PolG mutator (D257A) mouse, which has previously been shown to display substantially increased mutations and deletions in mtDNA^26^. In the RMC assay, we amplified two different regions of the mtDNA, one that spans a region containing a *Taq*I restriction enzyme (RE) site (Taq634), and another that does not span a *Taq*I site (mtCont). Before PCR analysis, the mtDNA from all samples are completely digested with *Taq*I enzyme so that in WT mtDNA, there is expected to be very little amplification with the Taq634 primer set (because the DNA template is destroyed by *Taq*I digestion). When random mutations occur throughout the mtDNA, some mutations will by chance occur randomly at the *Taq*I RE site rendering it resistant to degradation. Thus, in this setting amplification will occur. In models in which these random mutations are expected to occur more frequently (i.e. Global PolG Mutator or Muscle specific PolG^mut^), amplification across the Taq634 site should increase accordingly. This being the case, as expected we did indeed observe a higher level of amplification across the Taq634 site from mtDNA isolated from global PolG Mutator muscles (purple bar), compared to aged match control samples (**Figure 1E)**. Interestingly however, from our PolG^mut^ muscle we did not observe any difference in the mutation load of mtDNA, in samples from either short- or long-term muscle specific truncation of the PolG protein. This result was unexpected given we were deleting the proofreading domain of PolG, which thus prompted us to investigate other aspects of mtDNA integrity.

A further readout of mtDNA integrity that can be obtained from ddPCR using the above assay, is an estimation of deletion rate in the mtDNA. Because we accurately load equal amounts of mtDNA in each ddPCR reaction, when amplifying with the mtCont primer set we can determine if the primers amplify equally between genotypes. With equal amounts of mtDNA template present, amplification should be equivalent between samples, and any reductions in this amplification would indicate a loss of template, or “deletion” of the target region. When we performed this assay on global PolG Mutator mtDNA, we noted that there was a minor reduction in the amplification of mtCont amplicons between WT and Mutator muscles (purple bars), consistent with the previously described presence of some deletions in this model (**Figure 1F**). However, when we performed the same analysis in our PolG^mut^ muscles (red bars), we observed a substantial reduction in amplification with the mtCont primers compared to PolG^cont^ muscle, particularly in the short-term cohort, suggesting that more than half of the mtDNA molecules in the PolG^mut^ muscles harboured deletions. Consistent with this finding, when we used conventional PCR to amplify a 10kb section of mtDNA from these same samples which is prone to deletion, we observed amplicons of many various sizes (smears) in PolG^mut^ mtDNA, compared to a mostly single consistent amplicon of 10kb in PolG^cont^ mtDNA (**Supplemental Figure 1G**). Together with the ddPCR data, these findings provide strong evidence that mtDNA from PolG^mut^ muscles are truncated with the presence of mtDNA deletions, confirming a functional molecular consequence related to our PolG model.

Finally, an additional function of PolG relates to its role in facilitating mtDNA transcription and providing template for the transcriptional machinery to induce gene expression from the mtDNA. In order to demonstrate that mtDNA transcription was not impaired in our model, we performed a bespoke RNA-sequencing analysis to assess global mitochondrial transcription, as previously described ^27^. Our data demonstrate that contrary to a reduction in mtDNA transcription, we in fact observe a robust global increase in RNA expression across both the positive and negative mtDNA strands in PolG^mut^ muscles compared to control (range from 0.5-3 Log_2_ fold increase) (**Figure 1G**). This is more clearly demonstrated using a heat map of expression of the protein coding RNAs from the mtDNA, which were all significantly increased in PolG^mut^ muscles (**Figure 1H**). Interestingly, the 7S RNA (mt-7s) was also robustly increased, which is a key rate limiting step of mitochondrial gene transcription (**Figure 1H**). Thus, it does not appear that our PolG mutant leads to any impairment in mtDNA transcription, however instead, is inducing an as yet unexplained increase in mitochondrial transcription.

Thus, the overall characterisation and validation of our inducible muscle specific PolG^mut^ model confirms successful deletion of the targeted exonuclease domain in the mRNA, and expression of a detectable (presumably truncated) PolG protein. This truncated protein does not appear to substantially negatively impact the replication of mtDNA, mtDNA transcription, or proteins important in maintaining mtDNA integrity. It does however lead to an increased abundance of mtDNA deletions, but does not increase the presence of mtDNA point mutations.

### Health Impacts of Muscle Specific loss of PolG Exonuclease Activity

After validating that our model displayed the molecular characteristics of a muscle specific model of mtDNA deletions, we next investigated what impact this pathology had on whole body health, metabolism and physiology. To comprehensively study these outcomes in the short and long-term cohorts, we performed a range of phenotyping procedures on these animals. The first and possibly most striking observation noted in these animals, was a robust and rapid loss of body weight at approximately 4 months (16 weeks) post-induction (tamoxifen). This began to present in the short-term cohort (**Figure 2A)**, but was prominent and robust in mice from the long-term cohort (**Figure 2B)**. In these older mice, prevention of weight gain started from ∼4 months (16 weeks) post-tamoxifen, with substantial weight loss occurring over the subsequent 2 months, before stabilising and mostly persisting until the end of the study. This occurred only in the PolG^mut^ mice, and not in any of the other three control lines at any time point (**Supplemental Figure 2A & 2B)**. Using EchoMRI, we were able to demonstrate that mutant mice had only a very minor drop in lean mass at 5 months of age as shown by absolute lean mass (**Figure 2C**) and as a percent lean mass to body mass (**Supplemental Figure 2C)**. However, 12 month old mice demonstrated a significant reduction in lean mass from ∼20 weeks onwards, with the greatest reduction at 32 weeks (**Figure 2D** and **Supplemental Figure 2D**). A significant contributor to overall weight loss in these animals was reduced fat mass, which was significantly lower in the shorter term cohort from ∼ 15 weeks post-tamoxifen (**Figure 2E)**, where the longer term cohort of mice had lost approximately half of their fat mass by 32 weeks post-tamoxifen, compared to controls (**Figure 2F)**. These effects were observed even when expressed as a percentage of total body weight (**Supplemental Figure 2E & 2F)**, indicating a specific reduction in adiposity in these animals. The loss of lean mass did not impact isolated muscle weights in the short-term cohort (**Figure 2G**), however muscles were smaller in the longer term mutant mice with a significant reduction in muscle weights observed for *Tibialis anterior* (TA), *Extensor digitorum longus* (EDL) and *Gastrocnemius* at 12 months of age (**Figure 2H)**. Interestingly, *Soleus* muscle weights were not impacted in this model, which is curious given this muscle is almost exclusively red fibres, whereas other muscle types are either mixed or predominantly white fibres. Importantly, this loss in tissue weight was not observed for other organs in these animals including heart and liver for either the short- or long-term cohorts (**Figures 2I & 2J)**, indicating that these animals were not just smaller per se. The exception to this was epididymal fat mass (Epi), which was significantly lower in both young and aged cohorts, consistent with the above EchoMRI data indicating a loss in fat mass.

**Figure 2:**
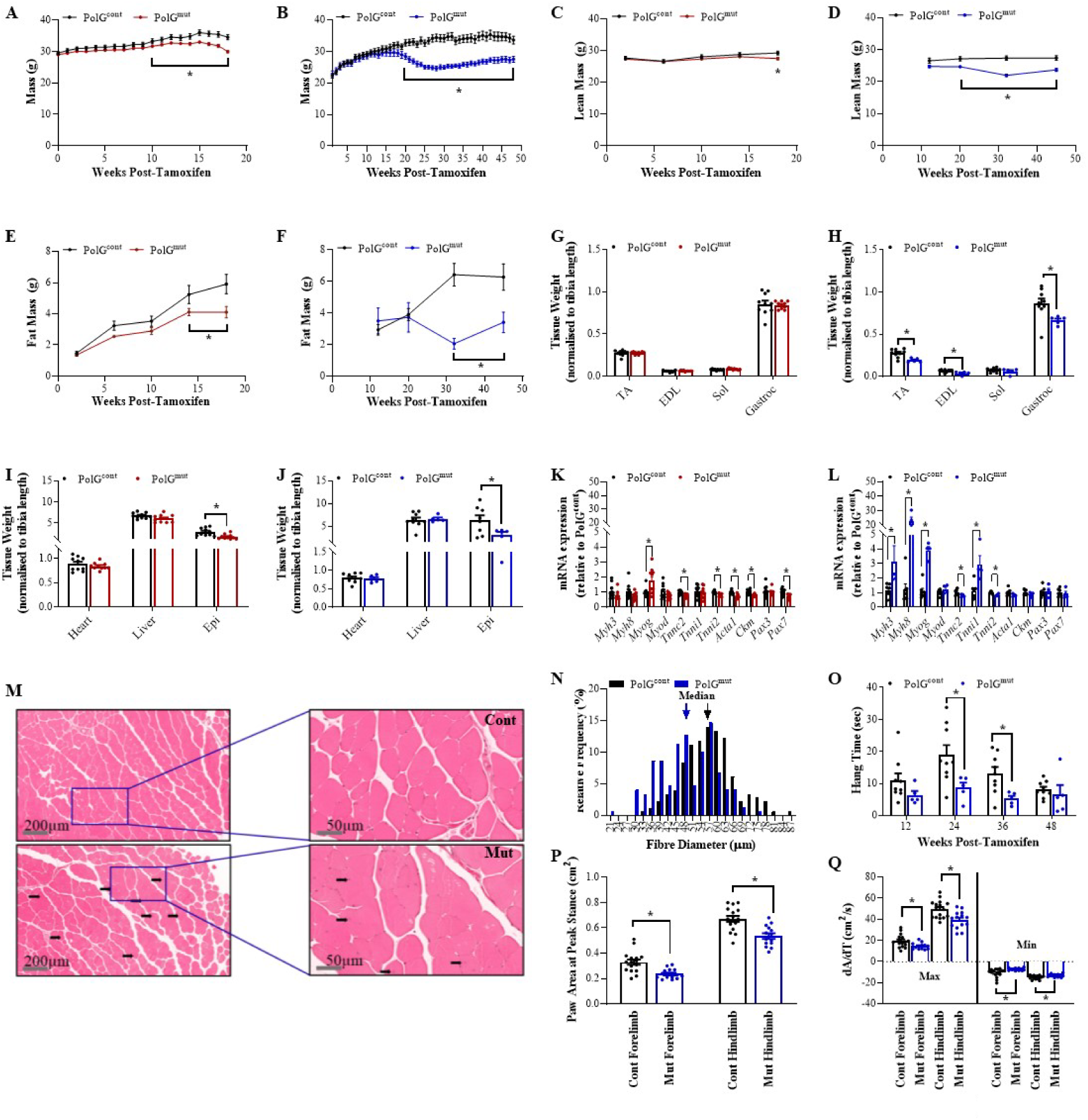
The Effects of Muscle Specific loss of PolG Exonuclease Activity on Muscle Function and Whole Body Parameters. Cohorts of PolG^cont^ and PolG^mut^ mice were studied over both short (4 months, red lines/bars) and long (12 months, blue lines/bars) time frames subsequent to the administration of tamoxifen. **(A)** Weekly body weights (mass) of mice in the short-term cohort and **(B)** weekly body weights (mass) of mice in the long-term cohorts. EchoMRI was used at various intervals throughout the studies to estimate **(C)** Lean mass in the short-term cohort, **(D)** Lean mass in the long-term cohort, **(E)** Fat mass in the short-term cohort and **(F)** Fat mass in the long-term cohort. **(G)** Individual muscle (TA, EDL, Sol, Gastroc) weights in the short-term cohort and **(H)** in the long-term cohort, **(I)** Heart, Liver and Epididymal white adipose tissue (Epi) weights in the short-term cohort and **(J)** in the long-term cohort. TA muscles were analysed for expression of genes associated with muscle growth and regeneration in the **(K)** short-term cohort and **(L)** in the long-term cohort. Sections of TA muscle were analysed using **(M)** H&E staining showing fibre size, integrity and nuclei position in control (Cont) and PolG^mut^ (Mut) mice in the long-term cohort. Boxes demonstrate magnified regions of each section (right panels), black arrows indicate centralised nuclei. **(N)** Quantification of fibre diameter distribution percentage in sections of TA muscle in the longer term cohorts, arrows indicate median fibre diameter. Functional assays to determine muscle strength and endurance were performed across the cohorts including **(O)** Bar graph plotting hang time (sec) duration as measured using the forearm hangwire test at various time points throughout the long-term cohorts. Data generated by DigiGait analyses in long-term mice demonstrating **(P)** Paw area at peak stance and **(Q)** Change of fore and hind paw area (dA/dT) during acceleration (min) and deceleration (max) when running. TA = Tibialis anterior, EDL = Extensor digitorum longus, Sol = Soleus, Gastroc = Gastrocnemius, Epi = Epididymal Fat Mass. Data are presented as mean ± SEM, n=5-10, * denotes p-value <0.05 between control and mutant samples as determined by repeated measures two-way ANOVA with correction for multiple comparisons (A-F) or unpaired, two-tailed Mann-Whitney t-tests (G-L & N-Q).

Given the prominent reduction in muscle weights in aged animals, we suspected that the exonuclease PolG mutant was likely inducing a muscle degenerative phenotype, consistent with many models in which mtDNA and mitochondrial function have been disrupted in myofibres^28–30^. Photographs taken of the animals at the end of their long term (12 month) experimental protocol provided additional evidence that the PolG^mut^ mice were smaller in stature and less muscular than control mice, with kyphosis (curved spine) being present in some animals, consistent with musculo-degenerative phenotypes (**White arrow; Supplemental Figure 2G**). To demonstrate degeneration of skeletal muscle more directly, we used qPCR to investigate transcriptional pathways in the skeletal muscle that would indicate altered expression of genes associated with myofibre degeneration and regeneration. Indeed, we observed a moderate reduction in mature muscle markers in the short-term cohort mice, with a reciprocal upregulation of Myogenin (*MyoG*), an early myogenesis marker

(**Figure 2K)**. The regenerative phenotype was more substantial in older mice, with robust upregulation of several markers of new fibre formation and myogenesis including *MyoG*, *Myh3*, *Myh8* and *Tnni1* (**Figure 2L)**. In support of these molecular changes, we also observed several fibres by histology that displayed centralised nuclei (**Black Arrows; Figure 2M)**, an indicator of new fibre formation, and an overall increase in the frequency of smaller diameter muscle fibres in mutant mice (**Figure 2N)**, all of which provide evidence that the PolG^mut^ muscle was undergoing bouts of degeneration and regeneration. Functionally, this remodelling and subsequent reduction in muscle mass, led to an overall loss of muscular endurance, as evidenced by a reduced latency to fall-time using the forearm hang wire test (**Figure 2O)**, which was most evident at 24 and 36 weeks post-tamoxifen. PolG^mut^ mice also exhibited postural and gait deficiencies compared with PolG^cont^ mice, as assessed by DigiGait analyses. This was indicated by reduced paw area (**Figure 2P**), and a reduced minimum and maximum dA/dT during walking at 33 weeks of age, which refers to limb loading time during the initial period of stance and the animal’s ability to propel itself into the next step (**Figure 2Q**). These findings are consistent with clinical observations often observed in human muscular degenerative conditions (cerebral palsy, muscular dystrophy) such as a hunched posture and a tendency to not walk appropriately on the flats of their feet/paws (i.e. toe walking). It is likely that compensation for these postural deficiencies has led to alterations in gait, which were observed in PolG^mut^ mice, particularly in the hind limbs (**Supplemental Figures 2H-2L**). Together, these analyses demonstrate that PolG^mut^ in skeletal muscle impacted on muscle form and function, and thus, posture and gait of these mice.

### Metabolic Impacts of Muscle Specific loss of PolG Exonuclease Activity

Our data above indicate that as PolG^mut^ mice age, their skeletal muscle is subjected to specific insults that lead to a degenerative phenotype and overall loss of muscle function and endurance. Such impacts would explain why muscle mass in these mice was smaller and frail, but are unlikely to explain why the animals had such profound changes in fat mass. Indeed, it would be expected that with a worsened muscle function and less muscle activity, that animals might have had a greater adiposity due to reduced energy expenditure. This is particularly confounding given that we did not detect any reductions in food intake in these animals in either the short- or long-term cohorts, which may have otherwise explained the loss of weight (**Supplemental Figure 2M & 2N)**. Therefore, subsequent investigations aimed at focusing on how this phenotype led to changes in fat mass and what, if any molecular underpinnings led to these systemic impacts on metabolism.

To understand this further, we investigated outcomes on whole body energy expenditure and glucose tolerance. Using the Promethion Metabolic Cage system it was apparent that there were no major differences in Respiratory Exchange Ratio (RER) in the shorter term cohorts (where body weights were mostly equivalent) (**Supplemental Figure 3A & 3B)**, but that long-term cohorts of PolG^mut^ mice demonstrated a minor (not significant) elevation in RER in the night period (**Supplemental Figure 3C & 3D)**. Similarly, 5 month old mice also demonstrated no difference in Energy Expenditure (EE) over 24 hours, although the expected increase in EE was observed in the active night period (**Supplemental Figure 3E & 3F)**. Notably, EE was significantly reduced in PolG^mut^ mice in the 12 month cohort of animals, which was consistent across both the day and night periods (**Supplemental Figure 3G & 3H)**, likely reflecting the reductions in body weight at this age. To test for this specifically, we performed ANCOVA analysis, where we observed no difference in EE in both young and older animals (**Supplemental Figure 3I & 3J)**, strongly suggesting that the EE differences observed in the older PolG^mut^ mice was indeed due to differences in body weight. Furthermore, we did not observe any difference in activity or movement, as X-Y-Z beam breaks across both cohorts and time points showed no major changes in these parameters (**Supplemental Figure 3K & 3L)**, although there was a trend for reduced movement in aged PolG^mut^.

With regards to glucose metabolism, we sought to investigate this by performing oral glucose tolerance tests (oGTT). Data from these analyses demonstrated that younger cohorts had no difference in their whole-body tolerance to a standardized oral glucose bolus, as indicated by an equivalent glucose excursion curve between the groups (**Supplemental Figure 3M)**, whereas in the older mice, the PolG^mut^ group had improved glucose excursions (**Supplemental Figure 3N)**. We also measured plasma insulin at the 0 and 15 minute time point post-glucose during the GTTs in both young and aged mice, and demonstrated that insulin was significantly reduced 15 minutes post-glucose in PolG^mut^ mice in both short and long term cohorts compared with PolG^cont^ (**Supplemental Figure 3O & 3P)**. Given this is a time point in the GTT at which insulin secretion is increased in response to rising plasma glucose (as observed in PolG^cont^ mice), a reduced level of plasma insulin might suggest improved insulin sensitivity in PolG^mut^ mice.

Overall, in the context of food intake and EE data above, these findings indicate that the large differences in fat mass between PolG^mut^ and PolG^cont^ mice, led to measurable improvements in whole body glucose tolerance and likely insulin sensitivity, which was most obvious in older mice. This is somewhat expected given that many previous studies have demonstrated that large reductions in fat mass result in similar phenotypic outcomes. However, what factors are driving this metabolic benefit, most notably weight loss in the PolG^mut^ mice, remains unknown at this point.

### Mitochondrial-centric Changes Induced by Muscle Specific loss of PolG Exonuclease Activity

With animal phenotyping data demonstrating robust changes in body weights, systemic metabolism and muscle weight/function, we were interested in determining the molecular changes that were occurring in the mitochondria of PolG^mut^ mice, which might explain such outcomes. We chose to analyse muscles in the short-term cohort of mice for these experiments, as these samples were collected at the critical point just prior to when the majority of weight loss occurred, allowing us to investigate the phenotype without major confounding consequences related to weight loss/muscle regeneration. To investigate these changes, we isolated intact mitochondria from quadriceps muscles of PolG^mut^ and PolG^cont^ mice and subjected them to proteomic analysis to obtain an unbiased quantitation of the mitochondrial proteome. Here, a tissue-centric label-free data-dependent acquisition proteome strategy was employed ^31^(n=8/group, ∼1300 proteins) quantifying 2275 proteins in total; 1319 proteins in the PolG^cont^ mitochondria, and 1353 proteins in PolG^mut^ mitochondria, which collectively covered 795/1140 of the known mitochondrial proteome (**Supplemental Figure 4A)** as previously described in MITOcarta 3.0 ^32^. Hierarchical clustering of differentially regulated proteins demonstrated that the proteins which were altered, were consistent amongst individual samples of each genotype, and in some instances were different in their z-score by up to 6 units (**Figure 3A**). Principle Component Analysis (PCA) demonstrated a strong influence of the two top components, which explained ∼30% (PC1) and ∼13% (PC2) of the variance respectively and was able to statistically separate the genotypes robustly based on these components (**Figure 3B**). Analysis and presentation of specific differentially expressed proteins (adj p<0.05) using a Volcano plot (**Figure 3C**), identified 68 proteins upregulated (red dots) in PolG^mut^ mitochondria, and 142 downregulated (blue dots). The top 20 most significantly regulated proteins revealed key members of pathways involved in ETC activity, mitochondrial ribosome function and cellular stress responses (**Supplemental Figure 4B**). Pathway enrichment analysis of differentially regulated proteins supported these observations, with a highly significant (q<1e^-30^) enrichment in downregulated pathways associated with “mitochondrial respiratory chain”, “mitochondrial electron transport” and “mitochondrial ATP synthesis” (**Figure 3D**). Regarding upregulated pathways, we observed less significant though broader enrichments, with the strongest being observed in pathways associated with alternative metabolism and metabolic processes (carboxylic acid, proline metabolism, amino acid catabolism), as well as protein import and cellular stress responses. As a way of confirming the strongest enrichments, we plotted two of the most regulated pathways using volcano plots to visualise the robustness of these changes, which demonstrated striking consistency in the downregulation of the vast majority of proteins associated with specific ETC complexes, including Complex I, III and IV (**Figure 3E**), and in proteins involved with mitochondrial ribosome/protein translation (**Figure 3F**).

**Figure 3:**
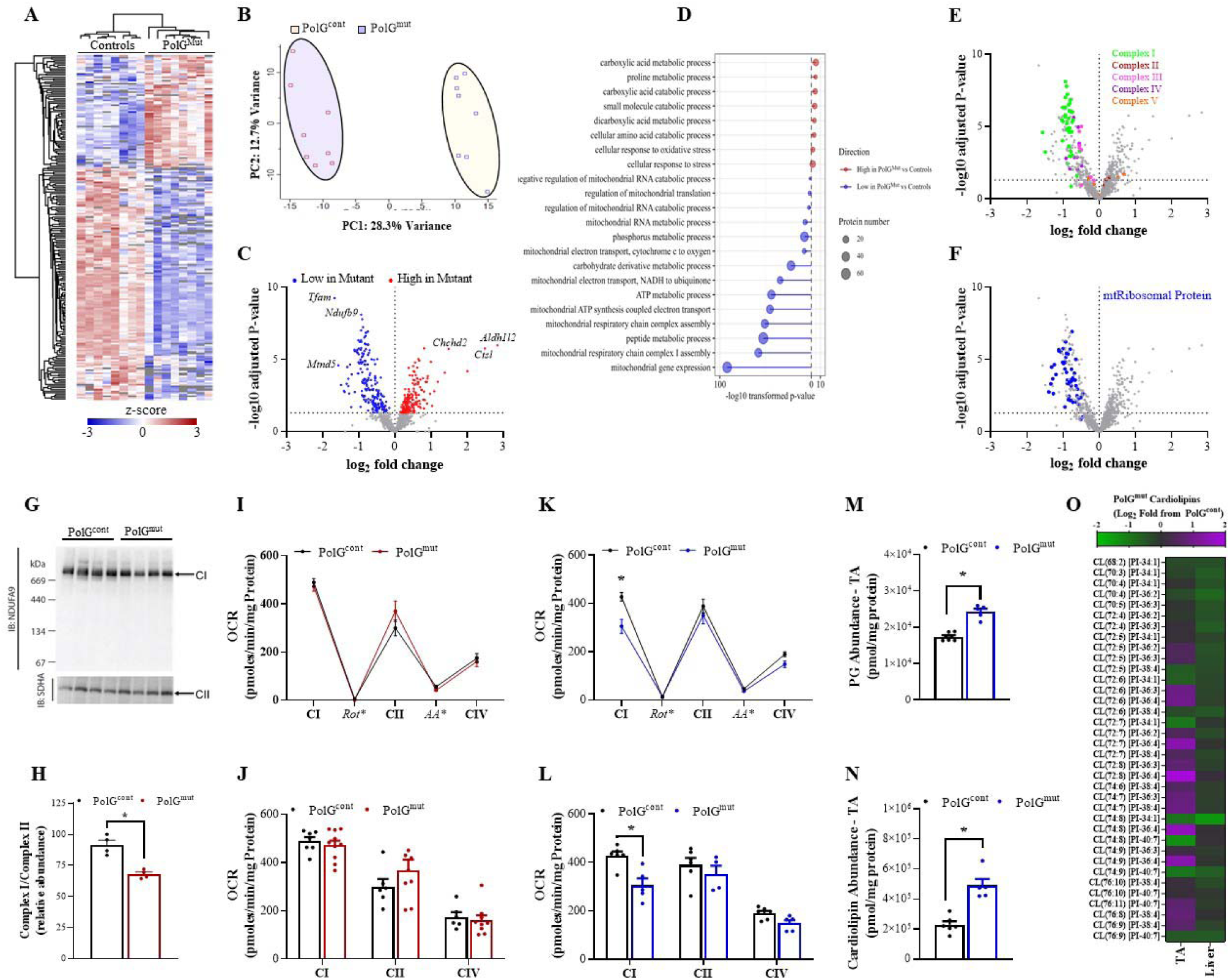
Mitochondrial Specific Changes Induced by Muscle Specific loss of PolG Exonuclease Activity. Unbiased assessment of protein abundance in mitochondria from skeletal muscle of PolG^cont^ and PolG^mut^ short term mice was analysed using proteomics. Computational analysis of the data was performed including **(A)** hierarchical clustering of protein abundance between PolG^cont^ and PolG^mut^, **(B)** Principal Component Analysis (PCA) of global changes in protein abundance between PolG^cont^ (open red squares) and PolG^mut^ (open blue squares) and, **(C)** Volcano plot depicting proteins that were significantly altered, (low-blue dots and high-red dots) in PolG^mut^ mitochondria compared to PolG^cont^ (grey dots indicates non-significantly altered proteins between groups). **(D)** Enrichment analysis of proteins significantly regulated in mitochondria between the genotypes, depicting upregulated pathways in red, and downregulated pathways in blue. Size of the circle indicates the number of proteins that contribute to the cluster. **(E)** Volcano plot of proteins, with the individual proteins of the electron Transport Chain (Complex I-V) highlighted (CI = green, CII = Red, CII = pink, CIV = purple, CV = orange) and **(F)** Volcano plot highlighting proteins from the mitochondrial ribosome complex that were altered in mitochondria between the genotypes. Isolated mitochondria were also analysed for functional differences including **(G)** Blue Native PAGE gel with western blot for Complex I (NDUFA9) and Complex II (SDHA) formation and, **(H)** Quantitation of BN-PAGE blots. The Seahorse analyser was used on fresh mitochondria isolated from muscle to measure Oxygen Consumption Rate (OCR) for activity of Complex I (CI), Complex II (CII) and Complex IV (CIV) from **(I&J)** short-term cohort, and **(K&L)** long-term cohort. Lipidomics was performed in TAs of PolG^cont^ and PolG^mut^ from long-term cohorts demonstrating the **(M)** abundance of the cardiolipin pre-cursor lipid phosphatidylglycerol (PG), and **(N)** total cardiolipin (CL). **(O)** Abundance of individual CL species (fold change from PolG^cont^) in TA and Liver from PolG^cont^ and PolG^mut^ long-term mice (purple = increased, green = decreased in PolG^mut^). For proteomics n=6/group, and analysis of data were corrected using Benjamini-Hochberg FDR. Functional mitochondria data are presented as mean ± SEM, n=4-10, * denotes p-value <0.05 between control and mutant samples and as determined by unpaired, two-tailed Mann-Whitney t-tests.

The specific changes in individual ETC complexes was consistent with data from previous studies that have investigated models in which mtDNA machinery is disrupted ^33^. Complex I is a primary component of the electron transport system and its dysfunction is often linked to degenerative diseases ^34–38^, especially in post-mitotic tissues such as skeletal muscle. To confirm whether the fully formed mature Complex I enzyme, and not just individual protein components, were downregulated in PolG^mut^ mitochondria, we performed Blue Native PAGE (BN-PAGE) analysis in PolG^cont^ versus PolG^mut^ muscle. BN-PAGE data demonstrated that mature Complex I (∼980kDa) was indeed lower in PolG^mut^ muscle compared to PolG^cont^ muscle, and consistent with proteomics data, mature Complex II formation appeared to be unaffected (**Figure 3G**). Quantitation of the ratio of CI/CII confirmed the significant reduction in Complex I formation (**Figure 3H**), providing evidence that there may be reduced Complex I activity in PolG^mut^ skeletal muscle. In order to functionally test the activity of individual ETC components, we analysed isolated mitochondria from skeletal muscle of PolG^mut^ mice using the Seahorse XF96 Flux analyser, which allows for the determination of oxygen consumption rates for different components of the ETC. We performed these experiments on mitochondria from both short and long-term cohorts. These data demonstrate that there was no significant impairment in oxygen consumption rate (OCR) mediated by any ETC components in the young PolG^mut^ mitochondria (**Figure 3I & 3J**), but there was a reduction in OCR driven by Complex I in the mitochondria isolated from older PolG^mut^ mice (**Figure 3K & 3L**).

Consistent with changes in mitochondrial function and remodelling, we identified striking differences in cardiolipin (CL) abundance between PolG^cont^ and PolG^mut^ TA muscles using targeted lipidomics analysis. Specifically, we identified an overall increase in CL precursor lipids; phosphatidylglycerols (PG) (**Figure 3M**), as well as an increase in the total abundance of CLs in PolG^mut^ TA muscles (**Figure 3N**). CLs are a critical lipid found in the inner mitochondrial membrane that usually contain two Phosphatidic Acid (PA) groups (each containing two FAs acyl chains), and changes in their abundance are associated with major mitochondrial remodelling ^39^. These changes in CL abundance were not observed in a non-mutant tissue such as the liver in PolG^mut^ animals, or in that of most other classes of lipids in the PolG^mut^ liver and plasma (**Supplemental Figure 4C**). Interestingly, when we investigated the abundance of specific CL species in the PolG^mut^ TAs, we noted that the majority of CLs that were significantly increased, were those which contained at least one phosphatidic acid (PA) as 36:4, which likely represents a PA with two 18:2 containing acyl chain (suggestive of newly synthesised or unmodified CLs) (**Figure 3O)**, which were also some of the most abundant (**Supplemental Figure 4D**). Indeed, some CLs were also reduced in PolG^mut^ TA muscles, which were mostly CLs containing a 34:1 PA. These trends were not observed in the liver of PolG^mut^ mice where PolG function would be expected to be normal. What the relevance of this CL remodelling is to the phenotype remains unknown, but may reflect the alteration in activity of specific mitochondrial enzymes that are no longer able to modify CLs, or that there is increased generation of new CLs to compensate for dysfunction, consistent with what has been recently postulated in mitochondriopathies in mice ^1^.

Overall, these molecular and functional data from mitochondria isolated from PolG^mut^ muscles, indicate a striking molecular change including profound loss of ETC and mitoribosome components, that in younger mice precedes systemic weight loss and eventually leads to reduced Complex I activity and muscle wasting in older mice. Whilst consistent with what might be expected from mutations that intrinsically impact mtDNA integrity, the ensuing alterations in body weight in older cohorts of these mice remained unexpected and unexplained.

### Nuclear Transcriptional Responses to Deletions in mtDNA

With proteomic, functional and phenotypic readouts indicating an intrinsic defect in mitochondrial activity in PolG^mut^ mice, we were interested to understand how these impairments impacted globally on the muscle transcriptional response. Specifically, we sought to understand what compensatory and alternative pathways were induced in muscle cells to overcome such deleterious deficits, and to identify which of these pathways may explain the observed weight loss in PolG^mut^ animals. To achieve this, we performed bulk RNA-sequencing on TA muscles from the short-term cohort of PolG^mut^ mice, so as to identify those pathways preceding, and perhaps driving, systemic effects.

Sequencing analysis was performed on n=6 of both PolG^cont^ and PolG^mut^ TA muscles using a NOVAseq6000. Mapping of the identified sequences identified 11683 genes across all samples. Hierarchical clustering of differentially regulated genes demonstrated consistent differences amongst samples of each genotype, and that genotypes clustered within their representative groups at the highest level (**Figure 4A**). PCA demonstrated a strong influence of the two top components, which explained ∼22% (PC1) and ∼20% (PC2) of the variance respectively, and was able to statistically separate the genotypes robustly based on these components (**Figure 4B**). Analysis of specific genes that were significantly differentially regulated (q<0.05) identified 492 genes that were downregulated, and 676 genes upregulated in PolG^mut^ compared with control muscles (**Figure 4C**). The top 20 most significantly regulated genes revealed key players in pathways involved in amino acyl tRNA activity, autophagy and the integrated stress response (ISR) (**Supplemental Figure 5A**). Interestingly, of the top 50 most regulated genes, 15 out of 21 of the known cytoplasmic tAcyl-RNA synthetases were upregulated, providing a strong indication that the nucleus was attempting to compensate for an apparent defect in protein translation or amino acid abundance. Further analysis of the principal components revealed some granularity regarding pathways that were contributing to these components, which demonstrated the strongest influence from pathways including RNA processing, ribosome biogenesis and translation (PC1) (**Figure 4D**). Other components suggested contributions from the ETC and oxidative phosphorylation (PC3), and pathways important in cellular remodelling and adhesion (PC5). Pathway enrichment analysis of differentially expressed genes demonstrated significant enrichment in upregulated pathways consistent with those observed in mitochondrial proteomics, which were namely related to alternative energy metabolism and the response to cellular and unfolded protein stress (**Figure 4E**), with downregulated pathways associated with extracellular matrix reorganisation and pathways associated with injury repair (**Figure 4F**). With regards to the integrated stress response (ISR), more detailed analysis of the top 20 most regulated genes demonstrated that at least half of these have previously been identified as being involved in the ISR. Activation of the ISR was confirmed by Western blotting in a subset of PolG^cont^ and PolG^mut^ muscles, which demonstrated robust phosphorylation (Ser51) of eIF2α, a key initiating process in the activation of the ISR ^40,41^(**Supplemental Figure 5B-5D**). Of note, were two upregulated genes that encode for the peptide hormones FGF21 and GDF15, which are released due to mitochondrial stress ^42–45^. These proteins are of particular interest in our study because of their recently described impacts on systemic metabolism and weight loss, and indeed these proteins are currently being investigated as biomarkers for mitochondrial myopathies in humans ^44,45^, and as potential treatments for metabolic dysregulation ^46–50^.

**Figure 4:**
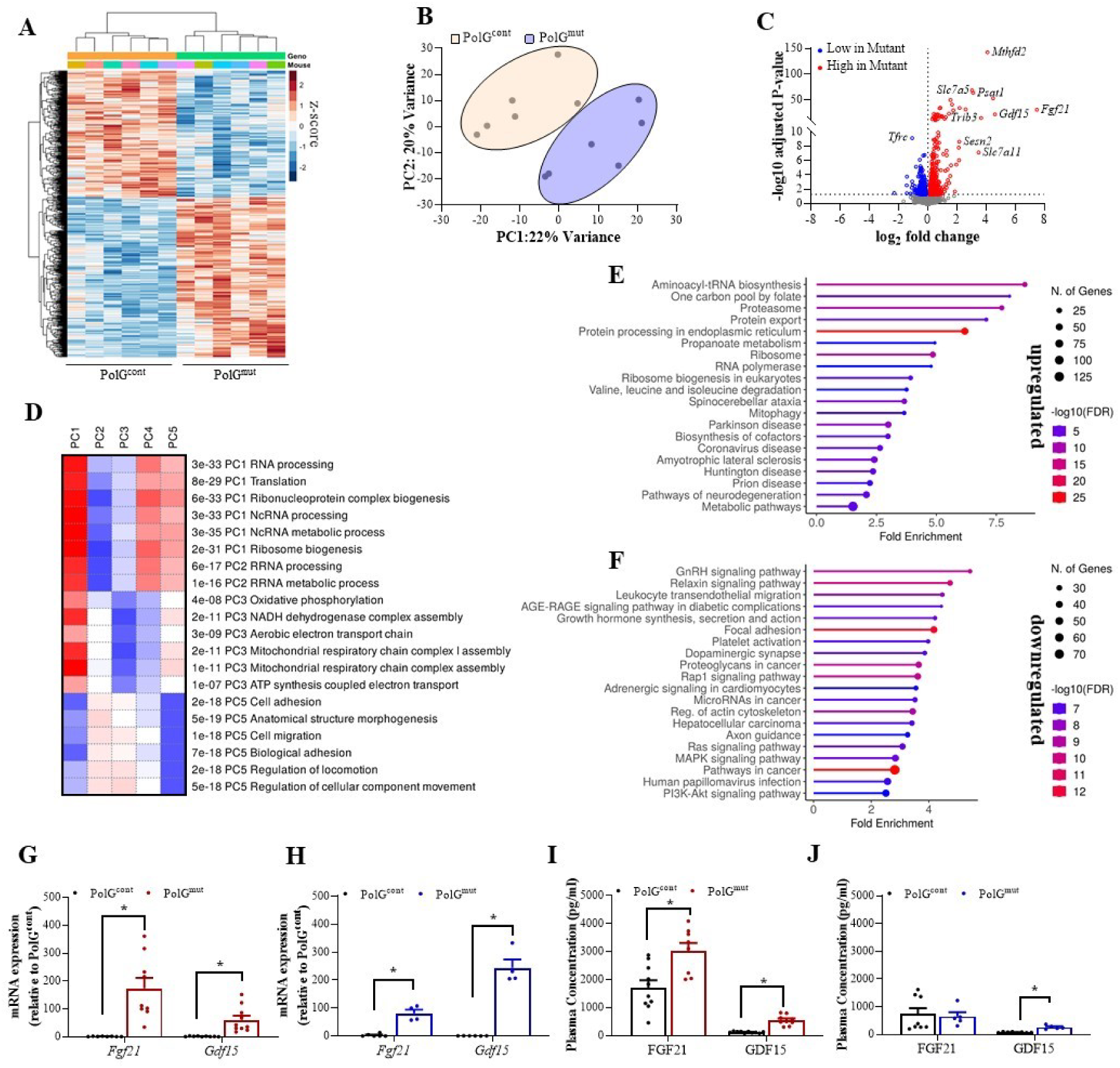
Transcriptional Response and Activation of the ISR in Response to Muscle Specific loss of PolG Exonuclease Activity. Unbiased assessment of RNA abundance was analysed from skeletal muscle of PolG^cont^ and PolG^mut^ short-term mice (4 months) using bulk RNA-sequencing. Computational analysis of the data was performed including **(A)** Hierarchical clustering of differentially expressed genes, **(B)** Principal Component Analysis (PCA) of global changes in RNA abundance between PolG^cont^ (beige shaded circle) and PolG^mut^ (blue shaded circle) and, **(C)** Volcano plot depicting genes that were upregulated (red dots) and downregulated (blue dots) in PolG^mut^ muscle compared to PolG^cont^ – grey dots indicate non-significantly altered genes between groups. **(D)** Specific identification of components driving global changes in principal component analysis (red = increased, blue = decreased, white = no change). Enrichment analysis (KEGG) on genes that were **(E)** significantly upregulated between the genotypes and **(F)** significantly downregulated between the genotypes. *Fgf21* and *Gdf15* mRNA expression as determined by qPCR in muscles from **(G)** short-term and **(H)** long-term PolG^cont^ and PolG^mut^ mice. Gene expression analysis was validated by ELISA for plasma abundance of FGF21 and GDF15 in **(I)** short-term and **(J)** long-term PolG^cont^ and PolG^mut^ mice. For RNA-seq n=6/group, and analysis of data were corrected using Benjamini-Hochberg FDR. Gene expression and ELISA data are presented as mean ± SEM, n=5-10, * denotes p-value <0.05 between control and mutant samples as determined by unpaired, two-tailed Mann-Whitney t-tests.

### Systemic Consequences of Muscle Specific activation of the Integrated Stress Response

With RNA-seq data demonstrating an approximate 180-fold and 25-fold increase in FGF21 and GDF15 respectively in PolG^mut^ muscle, we wanted to confirm these findings through targeted molecular assays. Firstly, we used qPCR to assess expression of these genes in skeletal muscle of all mice from each of the PolG^mut^ cohorts (short and long-term), to understand the temporal regulation of these peptide hormones. Consistently, we observed equivalent fold change in both FGF21 and GDF15 using qPCR as that measured by RNA-seq in the short-term cohort (**Figure 4G**). Interestingly however, we demonstrated in muscle from long-term PolG^mut^ mice, that FGF21 expression had reduced by about 50% (to ∼80-fold), whilst GDF15 expression continued to increase further to >200-fold above PolG^cont^ mice (**Figure 4H**). It is worth noting that we previously demonstrated that muscles at this time point were undergoing substantial regeneration (Figure 2), which may explain the reduction in FGF21; as damaged muscle would be replaced by new fibres that were PolG replete. However, GDF15 continued to increase, so whether this reflects a secondary chronic, or a feed forward effect of the ISR in aged fibres due to muscle regeneration remains unclear. To investigate this further, we examined whether the increased gene expression of these factors in muscle, resulted in increased release into the circulation. Indeed, we demonstrated that plasma levels of FGF21 were ∼2-fold higher in PolG^mut^ mice form the short-term cohort, whilst GDF15 was increased by several hundred fold (**Figure 4I**). This apparent discrepancy between gene expression and plasma levels was mostly due to plasma FGF21 being moderately abundant in young, healthy PolG^cont^ mice (1718.5pg/ml ± 251.8), and thus the increased contribution to plasma FGF21 induced by PolG^mut^ muscle was comparably minor. On the contrary, basal GDF15 abundance in healthy PolG^cont^ mice was comparatively lower (122.2pg/ml ± 9.1), and therefore the increased expression of GDF15 from muscle of PolG^mut^ had a major impact on plasma levels. These findings indicate that in healthy mice, a major proportion of FGF21 in the plasma of young mice arises from tissues other than skeletal muscle, whereas GDF15 is basally very low in the plasma of healthy mice, with the majority of GDF15 in PolG^mut^ presumably arising from stressed mitochondria in skeletal muscle. In mice from the long-term cohort, we noted that FGF21 abundance in the plasma of control mice was reduced by ∼3-fold (compared to younger mice), and indeed had reduced back to control levels in aged PolG^mut^ mice (**Figure 4J**). Plasma levels of GDF15 also reduced with age in PolG^cont^ mice, as did levels in PolG^mut^ mice, although the levels were still significantly elevated (∼3.5 Fold) compared to PolG^cont^ mice. Consistent with the observation that FGF21 levels were no longer elevated in aged PolG^mut^ mice, was the noticeable rebound effect in body weight and fat mass in aged PolG^mut^ mice (Supplemental Figures 2 & 2F). We surmise that these effects were related to regenerating fibres, and the subsequent reduction in FGF21 expression reflects an alleviation in skeletal muscle mitochondrial stress (as observed by reduced EIF2α phosphorylation in aged versus young animals; Supplemental Figure 5B & 5D). However, the enduring increase in GDF15 gene expression and sustained elevation in plasma GDF15 in PolG^mut^ mice is inconsistent with this effect on body composition, and appears independent of regenerating fibres. Therefore, we understand these findings to indicate that FGF21 was the primary driving factor impacting body weight and fat mass in these mice, and that GDF15 was not mediating further metabolic changes, unless compensatory mechanisms such as GDF15 resistance in peripheral tissues had prevented chronic activation of the GDF15 pathway (i.e. downregulation of the GDF15 receptor GFRAL).

### Upstream Pathways Leading to ISR activation in PolG^mut^ muscle

To decipher what leads to activation of the ISR and increased expression of FGF21 and GDF15 in PolG^mut^ skeletal muscle, we investigated several pathways of interest in our mutant muscle. Firstly, we did not observe any alterations in classic “energy sensing” pathways in the muscle of mutant mice, including the Akt, AMP-kinase and mTOR pathways (**Supplemental Figure 5E & 5F**). Indeed, we observed a significant reduction in phosphorylation of mTOR at both time points, indicating a suppression of signalling through this pathway which is inconsistent to what has been observed previously in the global Deletor (TWNK-KO) mouse ^42^. In the absence of any obvious metabolic pathway being involved in the molecular underpinnings of the response, we chose to focus on known activators of the ISR and Eif2α. We had already demonstrated that phosphorylation of Eif2α was increased in vivo in mouse PolG^mut^ muscle (Supplemental Figure 5B), and therefore we presumed that one of the four upstream kinases of Eif2α must also be activated. The four known kinases of Eif2α are PKR, PERK, HRI and GCN2, which are activated in response to specific cellular damage signals, some of which are known and others that are not ^51^. The two main kinases implicated in mitochondrial stress are GCN2 ^52^ and HRI ^53^, with PERK and PKR being more associated with ER stress and viral invasion ^54^. Therefore, we focussed on GCN2 and HRI pathways in our mutant mice.

GCN2 has been previously shown to be activated by mitochondrial ROS induced reductions in asparagine ^52^, and given we observed increased expression of asparagine synthase (Asns) in our model, we questioned whether this mechanism might be active in our mutants. However, western blots for phosphorylated GCN2 from skeletal muscle of both young and aged mice demonstrated no difference between PolG^cont^ and PolG^mut^ mice (**Supplemental Figure 5G & 5H**), suggesting that GCN2 is not the upstream pathway to ISR in our model. We subsequently shifted our focus to the HRI pathway, which was initially described in relation to its roles in iron metabolism ^55,56^, but more recent work has shown that mitochondrial ISR can be activated via this pathway in both an iron dependent ^57^ and independent ^58,59^ manner. Mechanistically, this is achieved via activation of the inner mitochondrial membrane protease OMA1, which cleaves the protein DELE1 in mitochondria, allowing cleaved DELE1 (sDELE1) to translocate from mitochondria to the cytosol where it binds and activates HRI ^58–60^. Interestingly, breaks in mtDNA have also recently been shown to activate the HRI/DELE1 pathway ^53^, although the precise mechanisms by which this occurs is not known. We show that in muscle from our mutant model, the abundance of HRI is increased (**Supplemental Figure 5I & 5J**), indicating that there is alteration in the homeostasis of this pathway. Furthermore, the shorter activated variant of the inner mitochondrial membrane protease OMA1, which is responsible for cleaving DELE1 and activating HRI, was also increased in our mutant model – again suggesting a disturbance to this pathway (**Supplemental Figure 5I & 5J**). These data suggest an alteration in HRI/DELE1 pathway, consistent with a role in the activation of the ISR in our model.

### Altered Folate Cycle Activity Underpins the PolG^mut^ ISR phenotype

With our data indicating a specific activation of the ISR in our mutant model, we were intrigued as to what the underlying initiators of this response might be. We reasoned that because many previous triggers of the mtISR have been shown to be metabolites, that alteration in abundance of a particular metabolite in our model may be the trigger. Therefore we performed a metabolomic screen on the muscle of PolG^mut^ and PolG^cont^ mice at both young and aged time points, to identify differentially abundant metabolites between genotypes. In the short and long term cohorts, we demonstrate that the metabolome is distinct enough to be separated by PCA (**Figure 5A & 5B**), but that only a handful of metabolites demonstrate a robust and significant (by FDR) change (**Figure 5C & 5D**). Enrichment analysis on the data of both young and aged metabolomes provided more insight, largely confirming findings from other analyses which identified major changes in alternative energy pathways (glycolysis, pentose phosphate pathway) and one carbon (1C) metabolism (**Figure 5E & 5F**). Whilst these enrichments were informative and confirmatory of previous investigations, they did not provide any insights into the temporal nature of the metabolomic changes. Therefore we plotted the individual metabolites of these enriched pathways across the two time points, to understand the temporal regulation in more detail (**Figure 5G**). These data highlight an interesting reversal of the metabolite profile with time, where for example glycolysis metabolites such as Glucose 6-phosphate and Fructose 6-phosphate were substantially reduced in young PolG^mut^ mice, but increased in aged mutant mice. Moreover, the pentose phosphate pathway was increased in young mutant animals, but decreased with age. Collectively these data highlight a major rewiring of energy metabolism over time in mutant mice, which indicate an initial attempt for muscle to divert resources through alternative mitochondrial energy pathways in young animals, but the cells eventually largely bypass mitochondrial energy metabolism in the aged animals to generate energy through multiple alternative pathways.

**Figure 5:**
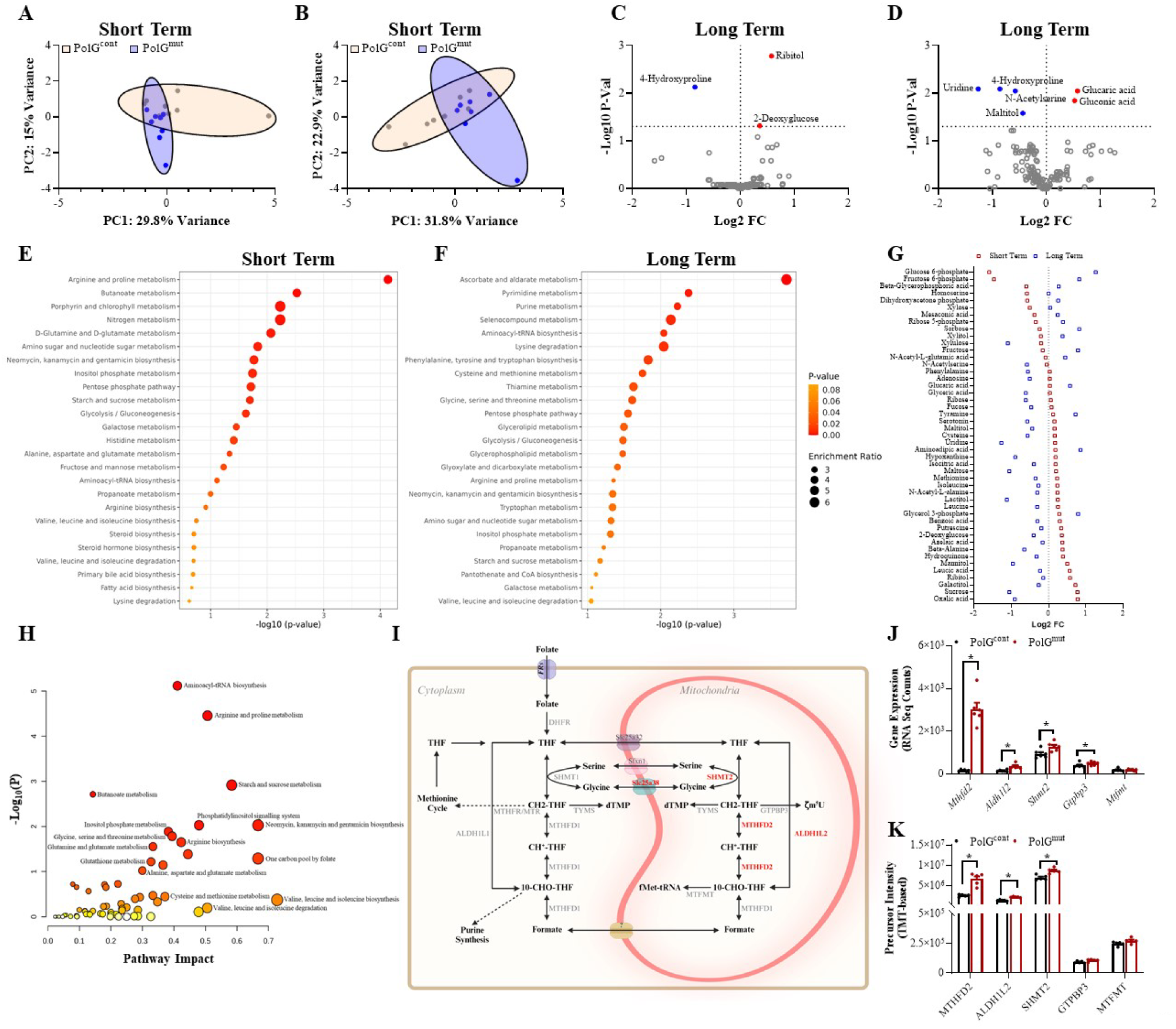
Metabolite Alterations in Muscle of PolG^mut^ Mice Mediated by Muscle Specific loss of PolG Exonuclease Activity. Metabolite abundance in skeletal mu scle from PolG^cont^ and PolG^mut^ mice was determined by metabolomics analysis. Computational analysis of the data was performed in the short and long-term cohorts including Principal Component Analysis (PCA) of global changes in skeletal muscle metabolite abundance in **(A)** short-term and **(B)** long term, between PolG^cont^ (biege shaded circle) and PolG^mut^ (blue shaded circle). Volcano plot depicting metabolites that were significantly upregulated (red dots) and downregulated (blue dots) in **(C)** short-term and **(D)** long-term PolG^mut^ muscle compared to PolG^cont^. **(E)** short-term and **(F)** long-term metabolic pathway enrichment analysis (MetaboAnalyst-KEGG) in order of significance with red gradient indicating P-Value and the size of the circle denoting the enrichment ratio. **(G)** Metabolites in PolG^mut^ skeletal muscle that had greater than 0.05-fold change (relative to PolG^cont^) between short and long-term analysis. **(H)** Integrated metabolic pathway analysis (MetaboAnalyst) of short-term skeletal muscle bulk RNA-sequencing and metabolomics data sets, with red gradient indicating p-value and circle size indicating pathway impact. **(I)** Schematic of the mammalian cytosolic/mitochondrial folate cycle (red text denotes upregulated proteins) in short-term cohort with **(J)** relevant folate cycle genes detected by RNA-sequencing and **(K)** proteins detected by mitochondrial proteomics. Metabalomics data was normalised by median and log transformed using MetaboAnalyst 5.0, n=8/group (except PolG^mut^ long-term cohort, n=7). For (J), RNA-seq n=6/group & (K) proteomics n=6/group and analysis of data were corrected using Benjamini-Hochberg FDR. * denotes p-value <0.05 between control and mutant samples.

A more encompassing method for understanding the underlying molecular causes of the mutant ISR phenotype, is to overlay and integrate omics outputs for additive benefit. Accordingly, we overlaid the RNA-seq and metabolite data from the short-term cohorts, in an attempt to capture consistent pathways that were being modulated prior to major adaptive changes in older mice. This approach highlighted many of the previous pathways identified by each analysis individually, but provided significant additional insights that were not previously captured by each method alone (**Figure 5H**). Specifically, we identified notable enrichments for butanoate metabolism, glycine/serine metabolism and one carbon (1C) Folate metabolism. In particular, we identify a major signal in the mutant muscle for enzymes and metabolites that support the synthesis of serine, and subsequently intermediates of the 1C folate cycle (**Figure 5I**). This is shown in Figure 5I in red text, which indicates proteins of the folate cycle that are significantly upregulated. Interestingly, serine is a critical co-factor for the enzyme SHMT2, which in the mitochondria catalyses the reversible conversion of serine and tetrahydrofolate (THF) to glycine and 5,10-methenyl-THF – the latter being a precursor to the formation of dUMP, dTMP (purine pathway) and to 10-formyl-THF, which is required for the biosynthesis of formyl-methionine (fMET). fMet is the tRNA responsible for initiating translation of many mitochondrial encoded RNAs ^61^, and its absence leads to stalling of mitochondrial translation. Furthermore, when revisiting our transcriptomic and proteomic data, we were intrigued to observe that two of the most upregulated genes/proteins in our omics datasets were ALDH1L2 and MTHFD2 (**Figure 5J & 5K**), both of which are critically important in regulating the abundance of 5,10-methenyl-THF. Our findings therefore point to a concerted effort for the mutant cells to increase the levels of THF intermediates, most likely 5,10-methenyl-THF. We suggest 5,10-methenyl-THF as there is no upregulation of TYMS or GTPBP3 (enzymes that increase dTMP and ζm5U respectively – indicating sufficient levels of 5,10-methylene-THF), and no increase in MTFMT or MTHFD1 (enzymes that increase fMet and formate respectively – indicating sufficient levels of 10-formyl-THF). Such evidence suggests that there is either an insufficient level of 5,10-methenyl-THF in mutant muscle, or an increased demand for this substrate.

Regarding the possibility for insufficient levels of 5,10-methenyl-THF, we identified above that there appeared to be a specific shuttle of resources towards increased serine synthesis, which likely reflects the cells attempt to increase activity of SHMT2 and thus synthesis of 5,10-methyleneTHF. This is demonstrated at both the RNA and protein levels for SHMT2 (**Figure 5J & 5K**). Yet this does not appear to be sufficient to remedy the pathologies present in our mutant muscle. Insufficient levels of 5,10-methenyl-THF would also be consistent with a block in mitochondrial protein translation (via flow on effects to 10-formyl-THF and subsequently fMet), which is strongly supported by our data demonstrating reductions in mitochondrial protein abundance, despite an upto 3-fold increased mitochondrial RNA transcription. Thus, our findings propose that reduction in the abundance of folate cycle intermediates is a contributor to mitochondrial dysfunction in this model, and may be a critical component leading to activation of the ISR.

In summary, we have generated the first inducible, tissue-specific PolG exonuclease deficient mouse model, which recapitulates many aspects of previous preclinical models that demonstrate decrements in PolG activity and mtDNA integrity, and importantly allows tissue specific investigations into key aspects of the condition. This includes an increased rate of mtDNA deletions and promotion of a degenerative phenotype in affected tissues. In this current study the tissue of interest was skeletal muscle, where 12 month old homozygous muscle specific mutant mice present with a musculo-degenerative phenotype that is preceded by a robust activation of the ISR. The ISR was activated alongside an apparent reduction in abundance of folate cycle intermediates, which may be a critical aspect of the subsequent activation of the ISR, and ultimately increased release of the peptide hormones FGF21 and GDF15 into the circulation. Collectively this promotes a systemic alteration in metabolism (in younger mice) and a resistance to increased body weight similar, as well as an aged related sarcopenia, all of which is consistent to that observed in the setting of cachexia (**Figure 6**).

**Figure 6:**
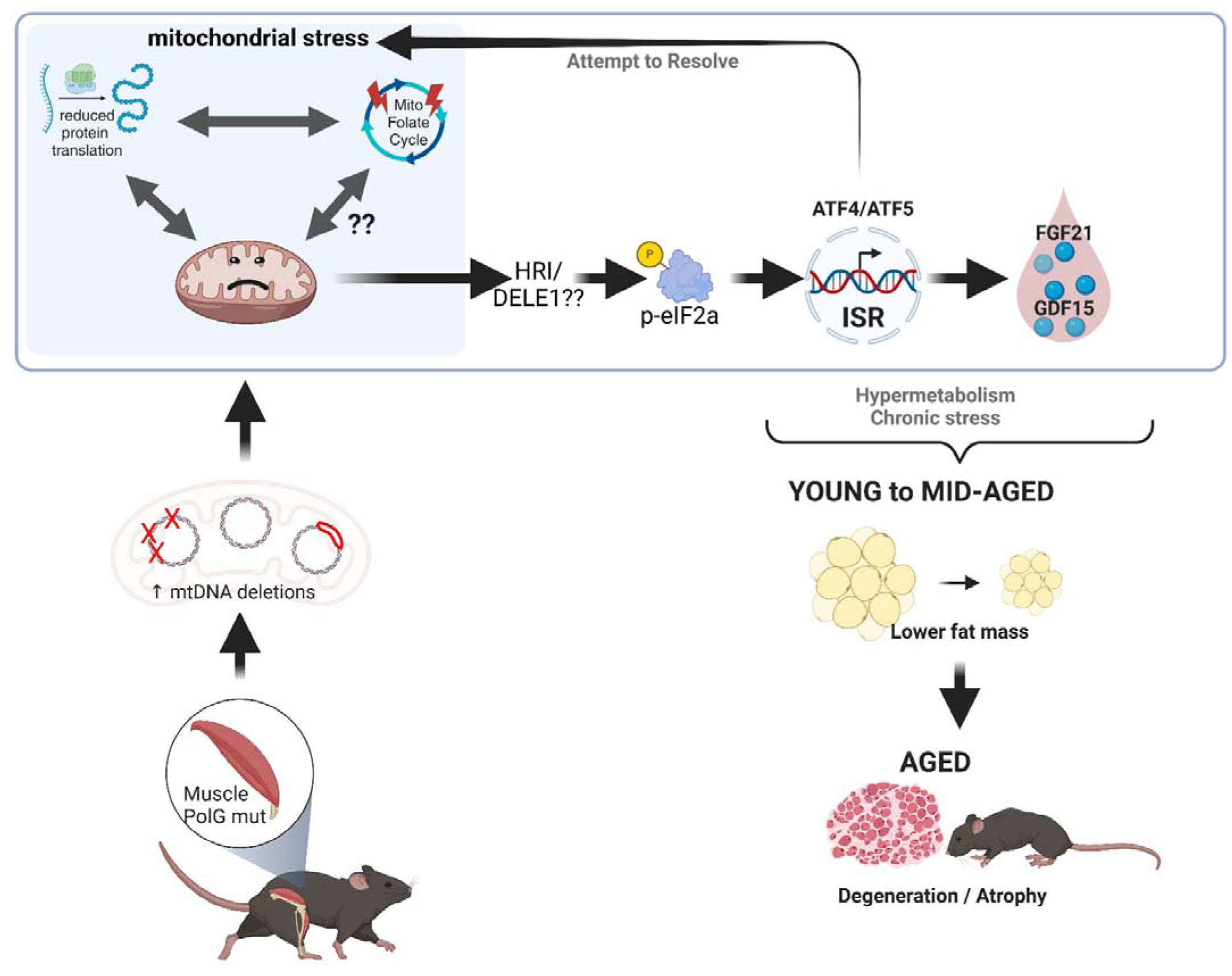
Schematic Summarising the Tissue-specific and Whole Body Impacts of Muscle Specific loss of PolG Exonuclease Activity. PolG Exonuclease deletion in skeletal muscle leads to an increased mtDNA deletion rate that with age promotes a degenerative phenotype specifically in the mutated tissue. This degeneration is preceded by a robust activation of the mitochondrial integrated stress response (ISR), driven in part by reduced protein translation and potentially reduced folate cycle metabolites. The activated ISR leads to an increased release of the peptide hormones FGF21 and GDF15 into the circulation. These series of events alter systemic metabolism that profoundly impact the accretion of fat mass, overall mirroring phenotypes consistent with cachexia.

## DISCUSSION

Herein we confirm and validate the generation of an inducible, tissue-specific mouse model of PolG exonuclease insufficiency, which importantly appears to retain all functions of PolG activity - other than the expected loss of mtDNA proofreading capacity (the specific function regulated by the exonuclease domain). These engineered defects in the PolG enzyme lead to an increase in the abundance of mtDNA deletions and mitochondrial heteroplasmy, which subsequently drives mitochondrial stress responses, perturbations and tissue pathology. We believe this is the first example of a conditional and temporal PolG mutator mouse model, which we anticipate will be a valuable model for the study of one of the most common inherited mitochondriopathies in humans. Moreover, using this unique model in a muscle specific context we have been able to reveal critical new knowledge in pathways associated with activation of the mitochondrial stress response – specifically that reduced abundance of specific folate derivatives may be important players in the ensuing activation of the integrated stress response.

In this study we investigated the short (4 month) and long (12 month) term impacts of post-developmental homozygous loss of PolG exonuclease activity specifically in skeletal muscle. It is worth emphasising that this is a post-developmental model, which was induced during adolescence and therefore all phenotypic changes are attributable to responses in mature muscle. Despite the post-developmental nature of this protocol, we still observed faithful recapitulation of key aspects of the global PolG mutator model, namely muscle degeneration and wasting. Whilst the animals demonstrated characteristics of skeletal myopathy, the animals did not display evidence of significant ill health up to 1 year post-deletion, with an observed mortality of 0% at this time point. This allowed us to study the whole-body impacts of this phenotype in the absence of complicating morbidities including neurodegeneration and heart failure – two common observations in the global mutator models ^23,62^. Unexpectedly we also observed a striking reduction in body weight at approximately 5 months post-mutation, driven at first by a loss of fat mass that with age also resulted in reduced lean mass, which reliably developed in all cohorts studied.

In depth molecular characterisation using proteomics, transcriptomics, lipidomics and metabolomics revealed that mitochondria from these animals had specific reductions in components of the electron transport chain (ETC) and mitochondrial ribosomal machinery. These defects were associated with a robust nuclear activation of stress signalling cascades, which did not include the type 1 interferon response, which has previously been shown to be important in global mutator models ^63^, but instead involved the mitochondrial unfolded protein response (mtUPR), integrated stress response (ISR) and alterations in alternative metabolism pathways. These classic hallmarks have been described in other models of “mitochondrial myopathy” and are collectively coined the integrated mitochondrial stress response (mtISR or ISR^mt^) ^42^. Whilst the mtISR likely elicited many cellular outcomes, the most prominent in our model was the elevation of muscle derived expression of the peptide hormones FGF21 and GDF15, and their subsequent release into the circulation. Given the large amount of published work and ongoing interest in these peptide hormones, we propose that it is the peripheral response to elevations in plasma FGF21 and GDF15 that likely explains the observed reductions in fat mass in these animals. This being the case however, we were surprised that this model did not demonstrate more striking alterations to metabolic readouts. Given the robust and chronic elevation in plasma FGF21 and GDF15, we expected to observe major differences in glucose/insulin tolerance, and energy expenditure (EE). Many recent papers have demonstrated that exogenous delivery of FGF21 and GDF15 promote substantial metabolic benefit to both rodents and humans via effects on food intake, EE, weight loss and improved glucose tolerance ^32,46–48,50^. Indeed, a very recent study also demonstrated that chronic exogenous delivery of GDF15 led to changes in body weight driven not only by reduced food intake, but in concert with a pathway in skeletal muscle that prevents the normal suppression of skeletal muscle energy expenditure ^64^. Whilst we observed robust reductions in fat mass and body mass, we were unable to attribute this weight loss to any major detectable differences in food intake or EE. This being said, we did observe a major difference in raw EE, but this was almost completely due to the differences in body weight, and once adjusted via ANCOVA, no significant difference was shown. Therefore, it is likely that small but prolonged changes in food intake and EE are likely responsible for the phenotype observed, that is below the level of detection with the methods we used. Consistent with an improved metabolic status, the glucose excursion curves following GTT were improved in older mice (but not younger mice), when there were much greater reductions in fat mass. This was accompanied by evidence for a likely improvement in insulin sensitivity, as demonstrated by a reduced plasma insulin following a glucose bolus during the GTT in both young and aged mice. In the context of the recent work by Wang et al demonstrating effects of GDF15 directly on muscle energy expenditure through calcium futile cycling, the body weight effects in our PolG model may relate to increased energy expenditure via this pathway. Whatever the mechanism, these mice in general had only a minor metabolic advantage, which might be useful information when assessing therapeutic options in humans with mitochondrial myopathy. Related to this, given that the mutant mice begun to regain weight in the latter stages of the aged experiment, our findings might indicate that chronic elevation of the peptide hormones FGF21 and GDF15, leads to a desensitisation of the periphery to these factors, highlighting important considerations for potential clinical trials planned with these targets.

Consistent with our data, recent literature describing similar intrinsic mitochondrial defects have also shown activation of these stress pathways, particularly in skeletal muscle. This includes models with impaired mitochondrial ribosome machinery ^65–67^, mitochondrial protein homeostasis ^68^, and mtDNA replication/polymerisation defects ^44,69^. Importantly however, our study is the first to demonstrate in a post-developmental PolG related model, robust and chronic increases in the mtISR, with induction of both FGF21 and GDF15 circulating hormones. There have indeed been select examples of increased plasma FGF21 being detected in the global PolG mutator model ^70^, however the literature regarding this has been inconsistent. Nevertheless, these findings collectively indicate that elevations in plasma FGF21 and GDF15 may be useful markers for the presence of mitochondrial disease, particularly those which manifest from respiratory chain defects, a proposal that has recently been tested and confirmed in small-scale human studies ^45,71^.

Why the ISR pathway and FGF21/GDF15 would be altered in the setting of skeletal muscle mitochondrial dysfunction remains incompletely understood, even though aspects of these pathways have been previously highlighted by several other groups. A teleological explanation for such an outcome is that specific mitochondrial insults elicit a starvation-like response in select tissues ^44^, in order to circumvent specific energy deficits. Indeed, FGF21 is increased in the liver in response to starvation ^72,73^, which potentially involves nutrient sensing pathways ^74^. In muscle, the mTOR pathway has been experimentally implicated in mitochondrial myopathy phenotypes together with activation of the mtISR, where global Twinkle-knock out mice (TWNK-KO/Deletor Mice) demonstrated increased activation of the mTOR pathway in muscle, whilst treatment with the mTOR inhibitor rapamycin, prevented activation of the muscle mtISR and reduced circulating FGF21 levels ^42^. Contrary to these findings, we did not observe activation of the mTOR pathway in our mutator mice, either in the short- or long-term cohorts, suggesting that this is not the mechanism by which the ISR is activated in our model.

Studies continue to investigate the underlying mechanisms relating to activation of the mtISR, with the majority of evidence suggesting that dysfunction in ETC activity is a likely contributor ^52,75–77^. Whilst this may be true in some instances, other studies have modelled defects in respiratory chain components in skeletal muscle, and rarely do these studies report weight loss associated with activation of the mtISR ^78^. Such observations have also been made in humans, where screening of plasma FGF21 and GDF15 in humans with known mitochondrial disease, demonstrated that FGF21 was almost exclusively elevated in individuals who had mitochondrial translation defects or mtDNA deletions, but not in other patients (including those with Complex I defects) ^79^. Thus, it is highly likely that a specific cascade of events must occur in skeletal muscle for the mtISR to be activated, and thus elicit this starvation-like response.

The hypothesis of a specific cascade of events activating the mtISR in skeletal muscle has been postulated and tested previously, particularly through the use of whole-body mouse models of mtDNA deletions (e.g. TWNK-KOs) ^44,80^. However as with existing PolG models, such global models cannot avoid complicating morbidities from other central and peripheral tissues, nor rule out defects acquired during development. For example, activation of the mTOR pathway in the Deletor mice (TWNK-KO) may be compensatory to signals received from other tissues, especially as we did not observe a similar activation in our model (where no compensation is likely). A recent study by Mick and colleagues systematically investigated alternative pathways that might lead to activation of the mtISR in cultured myoblasts and myotubes ^52^. This study also postulated that there are likely many pathways that can activate the mtISR, which are tissue, cell and context dependent. Ultimately, this group provided evidence for a disruption to amino acid metabolism that was a consistent component to their phenotype, with a specific reduction in Asparagine shown to activate the stress kinase GCN2 and subsequently the mtISR. Important insights from this and other studies hint to differential activation of the mtISR in various tissues, even though the same mitochondrial defect may be apparent. This appears to be consistent with work from other groups that implicate HRI, another Eif2α kinase, playing a role in activating the ISR in response to various stresses on the mitochondria including alterations to iron abundance and breaks in mtDNA. Our findings also suggest that HRI is the likely upstream kinase activating the ISR in our model, whilst also discounting a role for GCN2.

Another notable observation from our studies are the major reduction in mitochondrial ribosomal complexes and other specific protein complexes in the mitochondria, despite a significantly upregulated transcription of mtDNA encoded RNAs. Whilst this is a characteristic observation of mitochondrial stress, the explanations of what drives this phenomena are less well described. Specifically, our mitochondrial proteome data demonstrated a substantial depletion of proteins associated with mitochondrial ribosomal function. Conversely, we reveal a major upregulation of all the acyl tRNA synthetase (ARS) enzymes important for protein translation at cytoplasmic ribosomes. This likely highlights a feedback mechanism whereby a loss of protein translation in the mitochondria, subsequently signals for an increased protein translation in the cytoplasm (presumably of proteins that might otherwise be transported into mitochondria), and is a described outcome relating to activation of the ISR. It has been demonstrated previously by others that the ISR activates members of the ATF class of transcription factors, and that ATF5 specifically binds to Amino Acid starvation Response Elements (AARE) in the nuclear genome following mitochondrial stress, inducing transcription of genes important in one-carbon metabolism including *Mthfd2* and *Psat1* ^81^. We also observed elevation in these genes together with the many ARS genes (**Supplemental Figure 5**) consistent with an activation of the amino acid starvation response. Nevertheless, this increase in cytoplasmic translation activity was apparently unable to alleviate the defect, providing evidence that there was a potential deficiency in protein transport across the mitochondrial membrane, as has been proposed previously by others ^82^.

The loss of mitochondrial protein translation, and increase in one-carbon and amino acid related genes, fuelled our interests in understanding the underlying triggers for activation of the ISR. As mentioned above, whilst GCN2 and HRI have both been implicated in leading to activation of Eif2α following mitochondrial stress, the precise molecular triggers of this response are not clear. Our metabolomics analysis of the PolG mutant mice identified notable alterations in serine/glycine metabolism alongside major rewiring of pathways regulating the folate cycle. Work from others have also identified dysregulation of Serine metabolism as being an important component of cellular health particularly relating to mitochondrial dysfunction ^83,84^. Our integrative data sets encompassing transcriptome, proteome and metabolome identify a highly coordinated cellular response to promote the synthesis of folate derivatives, notably 5,10-methenyl-THF. This provides evidence of an underlying defect that might explain both the disruption to mitochondrial protein translation, and subsequent activation of the ISR. The reasons for why there is a deficiency in these metabolites in our model is not currently determinable from the current data, but proposes the exciting possibility that interventions which replenish these intermediates, might have therapeutic utility in the setting of various mitochondrial disorders.

In conclusion, our novel muscle specific PolG exonuclease mutant model provides several important insights into disease pathology driven by disruptions to mitochondrial integrity and the ISR in skeletal muscle. Notably, this was studied in the absence of complicating pathologies from other affected tissues, allowing for a longer term and more precise understanding of PolG function in skeletal muscle. Moving forward, our conditional model will provide the field with a novel opportunity to investigate mtDNA deletion-driven pathology in almost any tissue of interest, thus significantly advancing our knowledge of disease that is driven by such defects. These opportunities may lead to the identification of novel targets and biomarkers, as well as therapeutics that replenish 5,10-methenyl-THF, with potential relevance to the human condition.

## METHODS

### Animals

All animal experiments were approved by the Alfred Research Alliance (ARA) Animal Ethics committee (E/2030/2012/B), and performed in accordance with the ethical guidelines set out by the National Health and Medical Research Council (NHMRC) of Australia. Floxed PolG mutator mice (Polg^fl/fl^) were generated in collaboration with the Monash Genome Modification Platform using CRISPR to insert LoxP sites within the intron between exons 3&4, and exons 5&6 of the PolG1 gene (Figure 1A). Recombination was achieved using the Cre-Lox system by crossing the Polg^fl/fl^ mice with ACTA1-CreERT2 mice (C57BL/6J background, Jackson Laboratories, #025750) to generate Polg^fl/fl^-ACTA1-CreERT2^−/−(^Polg^fl/fl^) and Polg^fl/fl^ ACTA1-CreERT2^+/−^ (Polg^fl/fl-Cre^) animals. Male mice were bred and genotyped, before being randomly allocated into two separate groups each, totalling four groups altogether. Mice were aged to 6–8 weeks before one group of each genotype received oral gavage of Tamoxifen (80mg/kg) in sunflower oil for three consecutive days, whilst the other received sunflower oil alone. This resulted in four different groups: Polg^fl/fl+OIL^, Polg^fl/fl-Cre+OIL^, Polg^fl/fl+TAM^ (PolG^cont^), Polg^fl/fl-Cre+TAM^ (PolG^mut^). Mice were housed at 22°C on a 12 h light/dark cycle with access to food (chow: Specialty feeds, Australia) and water *ad libitum* with cages changed weekly. Several separate cohorts of PolG^mut^ mice were generated, which were either aged to 5 months (short-term) or to 12 months (long-term) post tamoxifen treatment, together with their respective controls. At the end of the studies, mice were fasted for 4–6 h and then anesthetized with a lethal dose of ketamine/xylazine before blood and tissues were collected, weighed and frozen for subsequent analysis.

### Glucose Tolerance Test (GTT)

An oral glucose tolerance test (oGTT) was performed at 46 weeks post-tamoxifen in in the long-term cohort and at 2, 6, 10 & 18 weeks post-tamoxifen in the short-term cohort, as previously described ^85,86^. Briefly, after a 5 h fast mice were administered a bolus of glucose (25% glucose solution) via oral gavage, at a dose of 2 g/kg calculated to lean mass as determined by EchoMRI. Blood glucose was measured at baseline before mice were gavaged, then subsequent measures of blood glucose were analysed at 15, 30, 45, 60, 90 & 120 min post-gavage using a glucometer (Accu Check Performa, Roche Diabetes Care). An insulin ELISA was conducted in conjunction with oGTT in the long-term cohort at 46 weeks post-tamoxifen and in the short-term cohort at 18 weeks post-tamoxifen. Plasma insulin quantification was determined using Mouse Ultrasensitive Insulin ELISA kit (ALPCO, USA) performed according to the manufacturer’s instructions using 5μl of plasma from 0 and 15 minute time points ^87^.

### EchoMRI

Body composition analysis, including lean mass (LM) and fat mass (FM), was analysed in living mice using the 4-in-1 NMR Body Composition Analyzer for Live Small Animals (EchoMRI LLC, Houston, TX, USA). Analysis was performed at designated periods throughout the study according to the recommendations of the manufacturer as previously described ^86–88^.

### Whole body energetics

Mice were placed in the Promethion High-Definition Behavioural Phenotyping System for Mice (Sable Systems International, North Las Vegas, NV, USA) at 34 weeks post-tamoxifen for long-term cohorts and at 8 weeks post-tamoxifen for short-term cohorts, for 3 consecutive days as previously described ^89^. Recordings for food intake, movement and respirometry, including energy expenditure (EE) and respiratory exchange ratio (RER), were collected over the final 24-h period following an initial acclimation period of 24 hours.

### Gait analysis (DigiGait)

Gait analysis was performed using the DigiGaitTM Treadmill System (Mouse Specifics Inc., USA) to assess different aspects of gait, posture, and motor function. DigiGait provides precise quantitative data measurements of gait parameters for each limb independently (left and right forelimbs and hindlimbs). Before each test mice were allowed to acclimatise in the Perspex chamber for 2 min. Digital images of animal paw placement from the ventral plane were then recorded through a clear treadmill that was running at a constant speed of 15 cm/s. Each mouse ran on the treadmill for approximately 2 minutes to ensure that a consistent 3-4 second period of uniform walking was recorded. For each mouse, videos were analysed using the DigiGaitTM Imaging and Analysis software v 12.2 (Mouse Specifics Inc., USA). Measurements were determined for each four limbs independently and reported as either separate limb values or combined and reported as forelimb and hindlimb values.

### Mitochondrial (mt)DNA to nuclear (n)DNA ratio

Mitochondrial content was determined by qPCR using a ratio of mtDNA to nDNA, as previously described ^89^. Briefly, *Tibialis anterior* muscle was homogenised in digestion buffer (100 mM NaCl, 10 mM Tris-HCl, 25 mM EDTA, 0.5% SDS, pH 8.0) and then incubated in Proteinase K (250U/mL) for 1 h at 55°C. Following this, total DNA was isolated using the phenol-chloroform extraction method. A qPCR reaction was then performed on 2ng of total DNA using a primer set that amplifies the mitochondrial gene *Mtco3*, and the nuclear gene *Sdha* (see **Table 1** for primer sequences). Estimated abundance of each gene was used to generate a ratio of mitochondrial to nuclear DNA (mtDNA/nDNA), and this ratio was compared between genotypes.

**Table 1:**
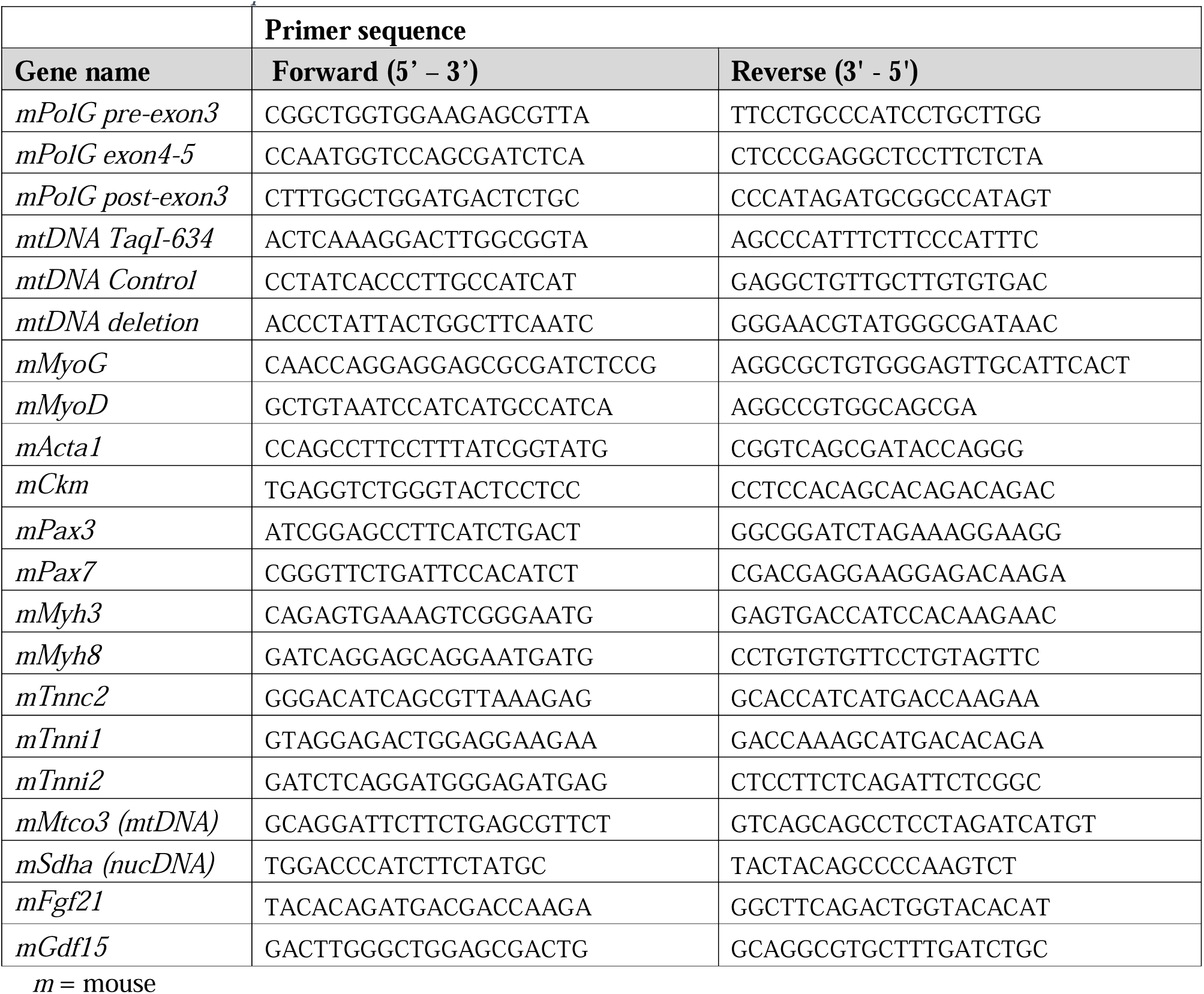
Primer sequences.

### Hangwire Strength Test

For latency-to-fall hangwire test, mice hang freely from their forepaws without the help of their hind legs or tail, on a wire suspended horizontally between two vertical beams 45 cm above a padded surface. Initially, mice underwent two days of training where mice were repeatedly placed on the wire for a total of 3 minutes per mouse. On day 3, mouse hangtime is recorded. Briefly, one test is completed when the animal lets go and drops onto the padded surface, which is repeated 3 times with a minimum 5 minute break between each test. The amount of time they were able to hang (latency-to-fall) was recorded and averaged over the three tests.

### Skeletal Muscle Transcriptomics

Bulk RNA sequencing was performed as previously described ^87^. RNA was extracted from *Tibialis anterior* (TA) muscles from each group using The Direct-zol RNA MiniPrep kit according to manufacturer’s instructions (Zymo Research). RNA quality for sequencing was determined using the high-sensitivity Agilent RNA ScreenTape Assay for 4200 Tape Station System (Agilent Technologies). Only samples with an RNA Integrity Number (RIN) score >8 were selected for subsequent analysis. DNA libraries were generated as per the manufacturer’s protocol using the NEBNext® Poly(A) mRNA Magnetic Isolation Module kit (New England Biolabs). 1μg total RNA per sample was inputted and adaptor ligation and PCR enrichment was performed using the NEBNext® Multiplex Oligos for Illumina® kit as per manufacturer’s instructions. Libraries for each of the 12 samples were individually bar coded using n6-barcodes and library quantity and quality were assessed as per manufacturer’s instructions using the Agilent D1000 ScreenTape Assay for 4200 TapeStation System. DNA libraries were pooled and then ran on a NovaSeq 6000 sequencer at the Centre of Genomic Medicine at the Alfred Hospital/Monash University. Reads were performed as paired-end with an average read length of 66 base pairs with a total of >400M reads collectively across all 12 samples, equating to approximately 30 million reads or more per sample.

### Transcriptomics Analysis and Gene Set Enrichment

RNA-sequencing data were processed and analysed similarly to that demonstrated previously^87^. Briefly, quality of raw sequence reads were assessed by FastQC. Based on the quality reports, read trimming was not required (Phred score > 35). Optical duplicates were removed using BBMap clumpify (https://github.com/BioInfoTools/BBMap) BBMap version 38.75. The processed reads were aligned to mm10 version of the mouse genome using STAR aligner. Mapped genes were counted using Feature Counts and lowly expressed genes were determined as those that had <1 count per million (CPM) in at least 8 samples, and removed. Trimmed Mean of M-values normalization (TMM) was performed to account for library size variation between samples. PCA was conducted with TMM normalised Log_2_ values to confirm separation of samples based on group, with no significant RNA concentration or read depth effect. Differential expression analysis was performed using DESeq2, ShinyGO.77, and iDEP.951 ^90^. Gene Set Enrichment Analysis of upregulated and downregulated gene sets was performed using ShinyGO.77 with Kyoto Encyclopedia of Genes and Genomes (KEGG) and Gene Ontology databases ^91^.

### Lipidomics

Lipidomics was analysed in plasma, TA muscle, and liver PBS homogenates as described previously in detail ^92^. Briefly, lipids were extracted from tissues/plasma using a 1:1 Butanol/Methanol extraction method, before application to ESI-MS/MS analysis ^92^. Quantification of lipids from MS analysis was performed using Mass Hunter Software (Agilent). Cardiolipins were annotated as their sum composition and the composition of the assigned product ion measured

### Metabolomics

Gas Chromatography Mass Spectrometry (GC-MS) polar metabolomic analysis was performed in skeletal muscle from young and aged mice (n=8/condition except long term PolG^mut^, n=7). Tissues were cryomilled in 600 µL of 3:1 methanol/water (containing 2 nmol of ^13^C_5_^15^N valine and ^13^C_6_ sorbitol as internal standards), using a Bertin Technologies Precellys bead mill coupled to a Cryolys cooling unit. The lysates were incubated at 4°C for 10 min with continuous agitation followed by centrifugation at 4°C for 10 min at 16000g, and supernatant transferred to fresh tubes. A 35µL aliquot of each sample was pooled to create the pooled biological quality control (PBQC). 35µL of each sample and the PBQC were transferred into HPLC inserts and evaporated at 30 °C to complete dryness in preparation for GC-MS analysis. Dried samples were derivatized online using the Shimadzu AOC6000 autosampler robot. Derivatisation was achieved by the addition of 25µL methoxyamine hydrochloride (30 mg/mL in pyridine) followed by shaking at 37°C for 2h. Samples were then derivatised with 25µL of N,O-bis (trimethylsilyl)trifluoroacetamide with trimethylchlorosilane (BSTFA with 1% TMCS) for 1h at 37°C. The sample was allowed to equilibrate at room temperature for 1 h before 1µL was injected onto the GC column using a hot needle technique. The GC-MS system used comprised of an AOC6000 autosampler, a 2030 Shimadzu gas chromatograph and a TQ8050NX triple quadrupole mass spectrometer (Shimadzu, Japan), with an electron ionization source (-70eV). Approximately 603 targets were collected using the Shimadzu Smart Metabolite Database, where each target comprised a quantifier MRM along with a qualifier MRM, which covers approximately 381 endogenous metabolites and multiple stable isotopically labeled internal standards. Resultant data was processed using Shimadzu LabSolutions Insight software, where peak integrations were visually validated and manually corrected where required. Enrichment and integration with transcriptomic data was performed using MetaboAnalyst 5.0 with Kyoto Encyclopedia of Genes and Genomes (KEGG).

### Quantitative PCR (qPCR)

RNA was isolated from tissues using RNAzol reagent and isopropanol precipitation. cDNA was generated from RNA using MMLV reverse transcriptase (Invitrogen) according to the manufacturer’s instructions. qPCR was performed on 10ng of cDNA using the SYBR-green method on a QuantStudio 7 Flex Real-Time PCR System, using primer sets outlined in **Table 1**. Primers were designed to span exon-exon junctions where possible, and were tested for specificity using BLAST (Basic Local Alignment Search Tool; National Centre for Biotechnology Information). Amplification of a single amplicon was estimated from melt curve analysis, ensuring an expected temperature dissociation profile was observed. Quantification of a given gene was determined by the relative mRNA level compared with control using the delta-CT method, which was calculated after normalisation to one of two housekeeping genes; *Ppia* or *Rplp0*.

### Mitochondrial Isolation

For isolation of mitochondria, *Quadricep* muscles were incubated and minced in ice-cold isolation buffer A (220mM Mannitol, 70mM Sucrose, 20mM HEPES, 2mM Tris-HCl, 1mM EDTA/EGTA, pH 7.2) + 0.4% (w/v) fatty acid free BSA and then homogenized with a glass dounce homogenizer. The homogenate was centrifuged at 650 x g for 5min at 4°C and the pellet was discarded, and the supernatant transferred to a fresh tube. The supernatant was repeatedly centrifuged at 650 x g for 5 minutes until very little material was pelleted. The supernatant was then transferred to a high-speed centrifuge tube and centrifuged at 10,000 x g for 5 minutes and the supernatant was discarded, and the crude mitochondrial pellet resuspended in isolation buffer A. Mitochondria were re-pelleted by centrifuging at 10,000 x g for 5 minutes, and the supernatant was discarded, and the pellet resuspended in 1ml of isolation buffer B (220mM Mannitol, 70mM Sucrose, 10mM Tris-HCl, 1mM EDTA, pH 7.2). The final pellet was collected by centrifuging at 10,000 x g for 5 minutes, discarding the supernatant and resuspending the pellet in either 1 x PBS for proteomic analysis or isolation buffer B.

### Random Mutation Capture Assay (RMCA) and Droplet Digital PCR (ddPCR)

The RMCA was performed similarly to that previously described^26^. Briefly, mitochondria were isolated from skeletal muscle as described above. Total DNA was then extracted using the phenol chloroform precipitation method. 50ng of mtDNA was digested for 6 hours with *TaqI* enzyme, with fresh enzyme added every hour before a final overnight incubation followed by inactivation at 80°C. 5ng of digested mtDNA was directly analysed on a BioRad QX200 Droplet Digital PCR System using primers that span the *TaqI* sensitive 634 site, where as 0.05ng of digested mtDNA was used for detection of the control, *TaqI* insensitive site (mtCont) (see **Table 1** for primer sequences). Assays were performed using EvaGreen reaction mix according to the manufacturer’s instructions using the following conditions: dissociation (95°C for 30s), annealing (61.8°C for 1min) and extension (72°C for 1.5min) for 50 cycles. Only samples where >10,000 droplets were quantified and included in subsequent analysis.

### Conventional PCR for mtDNA Deletions

To visualise mtDNA deletions we performed conventional PCR on isolated mtDNA from skeletal muscle, and resolved the amplicons on 0.8% agarose gels. Briefly, 10ng of mtDNA precipitated from skeletal muscle mitochondria (previously treated with DNAse I), was amplified using Phusion Polymerase according to the manufacturer’s instructions, using the “mtDNA deletion” primers outlined in **Table 1** (which flank the major arc of the mtDNA molecule; ∼10kb). The following conditions were used in the amplification reaction: dissociation (95°C for 30s), annealing (60°C for 30sec) and extension (72°C for 5min) for 40 cycles.

### SDS-PAGE and immunoblot

*Gastrocnemius* muscles were lysed in radio-immunoprecipitation assay (RIPA) buffer supplemented with protease and phosphatase inhibitors. Matched protein quantities were separated by SDS-PAGE and transferred to PVDF membranes. Membranes were blocked in 3% skim milk for 2 h and then incubated with primary antibody (see **Table 2** for details) overnight at 4°C. After incubation with primary antibodies, membranes were washed and probed with their respective HRP-conjugate secondary anti-mouse or anti-rabbit (BioRad) antibodies in 3% skim milk for 2h at room temperature, then visualised with enhanced chemiluminescent substrate (Pierce). Approximated molecular weights of proteins were determined from a co-resolved molecular weight standard (BioRad, #1610374). Image Lab Software (BioRad) was used to perform densitometry analyses, and all quantification results were normalized to their respective loading control or total protein.

**Table 2:** Antibodies for western blot.

| Antibody | raised in | size (kDa) | Company | Cat # |
| --- | --- | --- | --- | --- |
| Pan 14-3-3 | Mouse | 30 | Santa Cruz | sc1657 |
| Anti-Mouse HRP | Goat | - | Bio-Rad | 1706516 |
| Anti-Rabbit HRP | Goat | - | Bio-Rad | 1706515 |
| Phospho GCN2 Thr899 | Rabbit | 220 | CST | 94668 |
| Total GCN2 | Rabbit | 220 | CST | 3302 |
| Phospho mTOR Ser2448 | Rabbit | 289 | CST | 5536 |
| Phospho AMPK Thr172 | Rabbit | 63 | CST | 2535 |
| Phospho Akt Ser473 | Rabbit | 60 | CST | 4060 |
| Phospho eIF2a Ser51 | Rabbit | 37 | CST | 3597 |
| Total eIF2a | Rabbit | 37 | CST | 9722 |
| Pan Akt1/2 | Rabbit | 60 | CST | 4691 |
| NDUFA9 | Rabbit | 39 | In-house | - |
| EIF2AK1 (HRI) | Rabbit | 71 | MyBioSource | MBS2538144 |
| OMA1 | Mouse | 40/60 | Santa Cruz | sc-515788 |
| SDHA | Mouse | 71 | Abcam | ab14715 |

CST = Cell Signaling Technologies

### Blue Native-PAGE

Mitochondria from skeletal muscle were isolated by differential centrifugation as previously described ^38^. Protein concentrations were determined by bicinchoninic acid assay (BCA; Thermo Fisher Scientific). Mitochondria were aliquoted and frozen at −80°C until required. Blue Native (BN)-PAGE was performed as described previously ^38^. Briefly, mitochondria were solubilized in detergent buffer (20 mM Bis-Tris pH 7.0, 50 mM NaCl, 10% (w/v) glycerol) containing 1% digitonin for 10 minutes on ice, followed by centrifugation at 16,000 x g for 10 minutes. The clarified supernatant containing mitochondrial complexes was then added to one-tenth of the volume of 10x BN-PAGE loading dye (5% (w/v) Coomassie blue G250, 500 mM ε-amino n-caproic acid, 100 mM Bis-Tris pH 7.0) before loading onto the gel. Following electrophoresis, transfer onto PVDF membrane (Merck; IPVH00010) was performed using an Invitrogen Power Blotter System (Thermo Fisher) according to manufacturer’s instructions and membranes were incubated with primary and secondary antibodies (See **Table 2** for details). Bands were visualised with Clarity Western ECL chemiluminescent substrate (BioRad; 1705061) on a BioRad ChemiDoc XRS+ imaging system according to the manufacturer’s instructions. Densitometry was performed as described previously ^93^. Pixel intensity was measured in the BioRad software Image Lab (version 5.2.1) by taking the intensity of a region of interest. Background signal was subtracted by taking a region of the same size away from the region of interest. Signal intensities were then normalized to the intensity of the loading control and taken as a percentage of the highest sample intensity. Densitometry data was then analysed in GraphPad Prism (version 9.3.1).

### Plasma FGF21 & GDF15

Commercial ELISA kits for FGF21 (R&D Systems, #MF2100) and GDF15 (R&D Systems, #MGD150) were used to measure respective plasma concentration as per manufacturer’s instructions.

### Isolated Mitochondrial Respiration

Oxygen consumption rates were measured in isolated mitochondria from the quadricep muscle of mice, as previously described ^94^. To examine mitochondrial function and respiration, oxygen consumption in isolated mitochondrial preps was measured using an XFe-96 Seahorse Bioanalyzer (Agilent, USA). Protein was quantified in fresh isolated mitochondrial preps from quadricep muscle of mice using BCA (Pierce Reagent kit). A total of 3µg of isolated mitochondria in 180µl Mitochondrial Assay Solution (MAS) buffer (Sucrose 70mM, Mannitol 220mM, KH_2_PO_4_ 5mM, MgCl_2_ 5mM, HEPES 2mM, EGTA 1mM, BSA fatty acid free 0.2 %, pH 7.4 adjusted with KOH 1mol/L) was loaded into a 96 well seahorse cell culture plate (Seahorse XFe96 FluxPak) and centrifuged at 2,000 x g for 15 min to adhere the mitochondria to the bottom of the plate. To examine sequential electron flow through different complexes of the electron transport chain, an electron flow assay was performed. The electron flow assay was used to determine Complex I, Complex II and Complex IV mediated respiration with sequential injections of Rotenone 2µM, Succinate 10mM, Antimycin-A 4µM, and L-ascorbate 10mM + N,N,N′,N′-Tetramethyl-P-Phenylenediamine (TMPD) 100µM. For the electron flow assay, isolated mitochondria were adhered to 96 well plate in MAS media supplemented with Sodium Pyruvate 10mM, Malate 10mM and FCCP 4µM.

### Proteomics

Label free and tandem mass tag (TMT)^95^ nano liquid-chromatography/mass spectrometry (nLC/MS) proteomics was performed as previously described ^95,96^ with some modifications.

#### Sample homogenisation, protein reduction, alkylation and digestion

Semi-pure mitochondrial extracts were quantified by Bradford Protein Assay and 15µg of protein for each sample was reduced (10mM dithiothreitol, DTT) for 1 hr at 25°C and alkylated (20mM iodoacetamide) for 30 minutes at 25°C in the dark, before binding of protein to magnetic beads as previously described^97^. Magnetic bead slurry was prepared by mixing SpeedBeads™ magnetic carboxylate modified particles (Cytiva, 65152105050250, 45152105050250) at 1:1 (v:v) ratio, washing with MS-grade water and reconstituted to a final concentration of 100 µg/µL. The beads were added to the samples at 10:1 beads-to-protein ratio and ethanol added to a final concentration of 50% (v/v). Protein-bound magnetics beads were washed three times with 200 µL of 80% ethanol and reconstituted in 50 µL of 50 mM triethylamonium bicarbonat (TEAB) pH 8.0. Protein digestion was performed with Lysyl Endopeptidase (enzyme:substrate 1:100, 125-05061, Wako Pure Chemical Industries) and trypsin (enzyme:substrate 1:50, Promega V5113) overnight at 37°C with agitation (1,000 rpm). Peptide digests were collected from the supernatant, dried by vacuum centrifugation and stored at -80°C. For label-free strategy (n=8 per group), peptides were reconstituted in 0.07% trifluoroacetic acid and quantified by Fluorometric Peptide Assay (Thermo Fisher Scientific, 23290).

#### TMT peptide labelling and high-pH fractionation

For TMT-based labelling, anhydrous peptide digests were reconstituted in 50mM TEAB pH 8.5 and quantified by fluorometric peptide assay (23290, Thermo Fisher Scientific). Peptides were labelled with 11-plex TMT according to the manufacturer’s instructions with minor modifications (Thermo Fisher Scientific, A34808, lot UG287488/278919). In brief, each 11-plex experiment contained 10 different chemical tags for peptide labelling (i.e., groups labelled n=5; WT-MUT-WT-MUT-WT-MUT-WT-MUT-WT-MUT). The eleventh tag was used for generating reference channel made by a pooled peptide digest from all samples. A list of the sample labelling strategy is available in PRIDE proteomeXchange (PXD033029). Normalised peptide samples (15 µg) were labelled with 11 plex-TMT (A34808, Thermo Fisher Scientific) at 4:1 label-to-peptide ratio for 2 hr and quenched with 0.5% (v/v) hyoxylamine for 30 min at RT. Labelled peptide samples were acidified with 3% (v/v) formic acid (FA) and pooled into a new microtube. Pooled samples were further desalted using Sep-Pak tC18 96-well µElution (186002318, Waters). Desalted peptide elutions were lyophilised by vacuum-based speedVac for 1 hr and reconstituted in 25 mM ammonium formate, pH 10 for high pH reversed phase (RP) microscale fractionation. In-house X-RPS Stagetips were used for peptide binding and peptides eluted using acetonitrile (2-50%, v/v) in 25 mM ammonium formate, pH 10. A total of 16 fractions were pooled into 3 samples using alternating combinations and lyophilised by speedVac. Peptide samples were reconstituted in 0.07% triflouroacetic acid (TFA) and quantified using Colormetric peptide assay (23275, Thermo Fisher Scientific).

#### NanoLC and Mass Spectrometry

##### For label-free proteomics

Spectra acquired in data dependent acquisition on an Q Exactive HF-X benchtop Orbitrap mass spectrometer coupled to an UltiMate™ NCS-3500RS nano-HPLC (Thermo Fisher Scientific) as previously described^31^. Peptides (360 ng) were loaded (Acclaim PepMap100 C18 3µm beads with 100 Å pore-size, Thermo Fisher Scientific) and separated (1.9-µm particle size C18, 0.075 × 200 mm, Nikkyo Technos Co. Ltd) with a gradient of 2–28% acetonitrile containing 0.1% formic acid over 95 mins followed by 28-80% from 95-98 mins at 300 nL min-1 at 55°C (butterfly portfolio heater, Phoenix S&T). An MS1 scan was acquired from 350–1,650 m/z (60,000 resolution, 3 × 10^6^ automatic gain control (AGC), 128 msec injection time) followed by MS/MS data-dependent acquisition (top 25) with collision-induced dissociation and detection in the ion trap (30,000 resolution, 1×10^5^ AGC, 60 msec injection time, 28% normalized collision energy, 1.3 m/z quadrupole isolation width). Unassigned precursor ions charge states and slightly charged species (were rejected and peptide match disabled. Selected sequenced ions were dynamically excluded for 30 sec.). Data was acquired using Xcalibur software v4.5 (Thermo Fisher Scientific). For spectral library generation and high-pH fractionation, anhydrous peptide digests from MUT and WT/control (pooled) were individually reconstituted in 25 mM ammonium formate, pH 10 for high pH reversed phase (RP) microscale fractionation. In-house X-RPS Stagetips were used for peptide binding and peptides eluted using acetonitrile (2-50%, v/v) in 25 mM ammonium formate, pH 10. A total of 8 fractions were lyophilised by speedVac. Peptide samples were reconstituted in 0.07% trifluoracetic acid (TFA) analysed in single shot proteomics as described ^98^.

##### For TMT-based proteomics

TMT-labelled peptides were analysed on a Dionex 3500RS nanoUHPLC coupled to an Orbitrap Eclipse Tribrid mass spectrometer equipped with nanospray ion source in positive, data-dependent acquisition mode. Peptides were separated using an Acclaim Pepmap RSLC analytical column (Dionex-C18, 100 Å, 75 μm × 50 cm) with an Acclaim Pepmap nano-trap column (Dionex-C18, 100 Å, 75 μm × 2 cm), and a gradient of 3–80% (0-3 min, acetonitrile containing acetonitrile and 0.1% (v/v) formic acid over 240 min at 300 nl min-1 at 50 °C. Peptides were injected into the enrichment column at an isocratic flow of 5 μL/min of 2% (v/v) acetonitrile containing 0.1% (v/v) formic acid for 6 min, applied before the enrichment column was switched in-line with the analytical column. An MS1 scan was acquired from 375–1,500 m/z (120,000 resolution, isolation window of 0.7 Thomsons) in ‘top speed’ acquisition mode with 3 s cycle time on the 3 most intense precursor ions; ions with charge states of 2 to 7 were selected. AGC target was set to standard with auto maximum injection mode, followed by MS/MS data-dependent acquisition with high-field collision-induced dissociation and detection in the ion trap (5 ×104 AGC, 70-ms injection time, normalized collision energy of 38, 0.7 m/z quadrupole isolation width). Activation time of 35 ms and activation Q of 10. Analysis of fragment ions was carried out in the ion trap using the ‘Turbo’ speed scanning mode. Dynamic exclusion was activated for 30 s, with data was acquired using Xcalibur software. The mass spectrometry proteomics raw files and data analysis files have been deposited to the ProteomeXchange Consortium (http://proteomecentral.proteomexchange.org) via the PRIDE partner repository with identifier (label-free/library: PXD039029; TMT-label: PXD033029).

#### Proteomics: Data Processing and Bioinformatics

MS raw files were analysed using MaxQuant (v1.6.14) software^99^ and peptide lists were searched against musculus protein database (SwissProt (TaxID = 10090) (55,470; Jan 2021) including contaminants database with the Andromeda search engine. For LFQ-based analyses (in combination with spectral library), cysteine carbamidomethylation was selected as a fixed modification and N-terminal acetylation and methionine oxidations as variable modifications. Data was processed using trypsin/P and LysC as the proteolytic enzymes with up to 2 missed cleavage sites allowed. Further processing using match between runs (MBR) and label free quantification (LFQ) algorithm were employed. Peptides were identified with an initial precursor mass deviation of up to 7 ppm and a fragment mass deviation of 20 ppm with a minimum length of 7 amino acids. False discovery rate (FDR) was 0.01 for both the protein and peptide by searching against a reverse database. For TMT-based analyses, reporter ion MS2 (11plex TMT) settings were employed. Additional protein/peptide analysis was performed using Proteome Discoverer (v.2.5, Thermo Fisher Scientific) with the Sequest HT search engine in combination with the Percolator semi-supervised learning algorithm^100^. Fragment match spectral analysis in Proteome Discoverer was used to export sequence specific peptide spectra.

Contaminants, and reverse identification were excluded from further data analysis. High confident protein identification required more than one unique or razor peptides per protein group. Data analysis was performed using the Perseus (v1.6.14.0) package and R programming language ^98^. Protein intensities were log2 transformed and normalized using quantile normalization from R package preprocessCore. The histogram of the precursor intensity distribution and the boxplot of correlation covariance were visualized using R package ggplot2. Proteins with no missing values were subjected to downstream visualization and statistical analysis using Perseus of the MaxQuant computational platform ^101^. Proteins were subjected to PCA and student’s t-test. g:Profiler and Reactome databases were utilized for functional enrichment and network/pathway analysis, significance p<0.05.

### Histology

TA muscles were fixed in formalin and mounted in paraffin before 4µm sections were cut on a Leica microtome. All sections were stained with Hematoxylin & Eosin, and slide images were captured using an Olympus Slide scanner VS120 (Olympus, Japan) and viewed in the accompanying software (OlyVIA Build 13771, Olympus, Japan). Fibre diameter was assessed on imaged sections using Aperio Imagescope Software (Leica). Analysis was performed on at least 2 sections from each mouse, and at least 5 mice from each group. A minimum of 25 fibres were counted across 5 different fields of view per section.

### Statistical analyses

All data were expressed as mean ± standard error of the mean (SEM) unless otherwise stated. Normality was checked using Shapiro-Wilk normality tests. All statistical analyses in animal studies were analysed by repeated measures two-way ANOVA (for normally distributed data), other than where explicitly detailed. Lipidomics, tissue analysis and cell-based experiments were analysed by ANOVA with post hoc testing (Fishers LSD) where appropriate, or paired Student’s t test unless otherwise stated. Analyses were performed using PRISM9 software and a *p* value of *p* < 0.05 was considered statistically significant.

### Data inclusion and exclusion criteria

For animal experiments, phenotyping data points were excluded using the following pre-determined criteria: if the animal was unwell at the time of analysis, there were technical issues identified (such as failed beam breaks in Promethion), or data points were identified as outliers using Tukey’s Outlier Detection Method (1.5 IQR < Q1 or 1.5 IQR > Q3). If repeated data points from the same mouse/sample failed QC based on predetermined criteria or several data points were outliers as per Tukey’s rule, the entire animal was excluded from that given analysis (i.e., during glucose tolerance tests, indicating inappropriate gavage). For *in vivo* and *in vitro* tissue and molecular analyses, data points were only excluded if there was a technical failure (i.e., poor RNA quality such as RIN = <7, failed amplification in qPCR, failed injection in mass spectrometer) or the value was biologically improbable. This was performed in a blinded fashion (i.e., on grouped datasets before genotypes were known).

## ACKNOWLEDGMENTS

We acknowledge funding support from the Victorian State Government OIS program to Baker Heart & Diabetes Institute (BHDI). These studies were supported by funding from the BHDI Obesity & Lipid Program, as well as fellowship support to BGD and ACC from the National Heart Foundation of Australia (101789 and 105631, respectively), and to BGD from the NHMRC (Investigator Grant 2016530). LEF acknowledges support from the Mito Foundation and the NHMRC (Investigator Grant 2010149). We thank members of the MMA, LMCD, Metabolomics, and Molecular Proteomics laboratories at BHDI for their contributions. We also acknowledge support from Prof Mark A. Febbraio (BHDI & Monash Institute of Pharmaceutical Sciences) in generating the floxed mouse, and guidance and resources from Michael T Ryan (Monash University). We are grateful for platform support from the Monash Histology Platform, Department of Anatomy and Developmental Biology at Monash University, Natalia Carvajal and Andrew Perkins from the Centre of Genomic Medicine at the Alfred Hospital Monash University, The La Trobe University Proteomics and Metabolomics Platform, and the University of Melbourne Mass Spectrometry and Proteomics Facility. We also thank A/Prof Peter Crouch from the School of Biomedical Sciences, University of Melbourne, Australia for providing skeletal muscle samples from the global PolG Mutator Model. Various figure panels were created using BioRender.com, with all such figure content sublicensed accordingly.

## AUTHOR CONTRIBUTION

BGD conceived the study and co-designed the experiments with DCH, STB and EJK. BGD and DCH conceived and generated the PolG floxed mouse model. BGD, STB & EJK wrote the manuscript. STB, EJK, AZ, AWJ, SMW, YL & CY performed animal experiments and phenotyping. BGD, STB, EJK, SMW, CY, YL, KHL, HAF, KH, DWG, APN, SRC, DCH, PJM, LEF, DLR, AF, NED, DdS, DLR & ACC analysed data, processed tissue samples, and performed molecular and biochemical experiments. ACC, DWG, MI, MTR, AF & PJM provided reagents, experimental advice, and access to infrastructure and resources. All authors read and approved the manuscript.

## SUPPLEMENTAL FIGURES

**Supplemental Figure 1:**
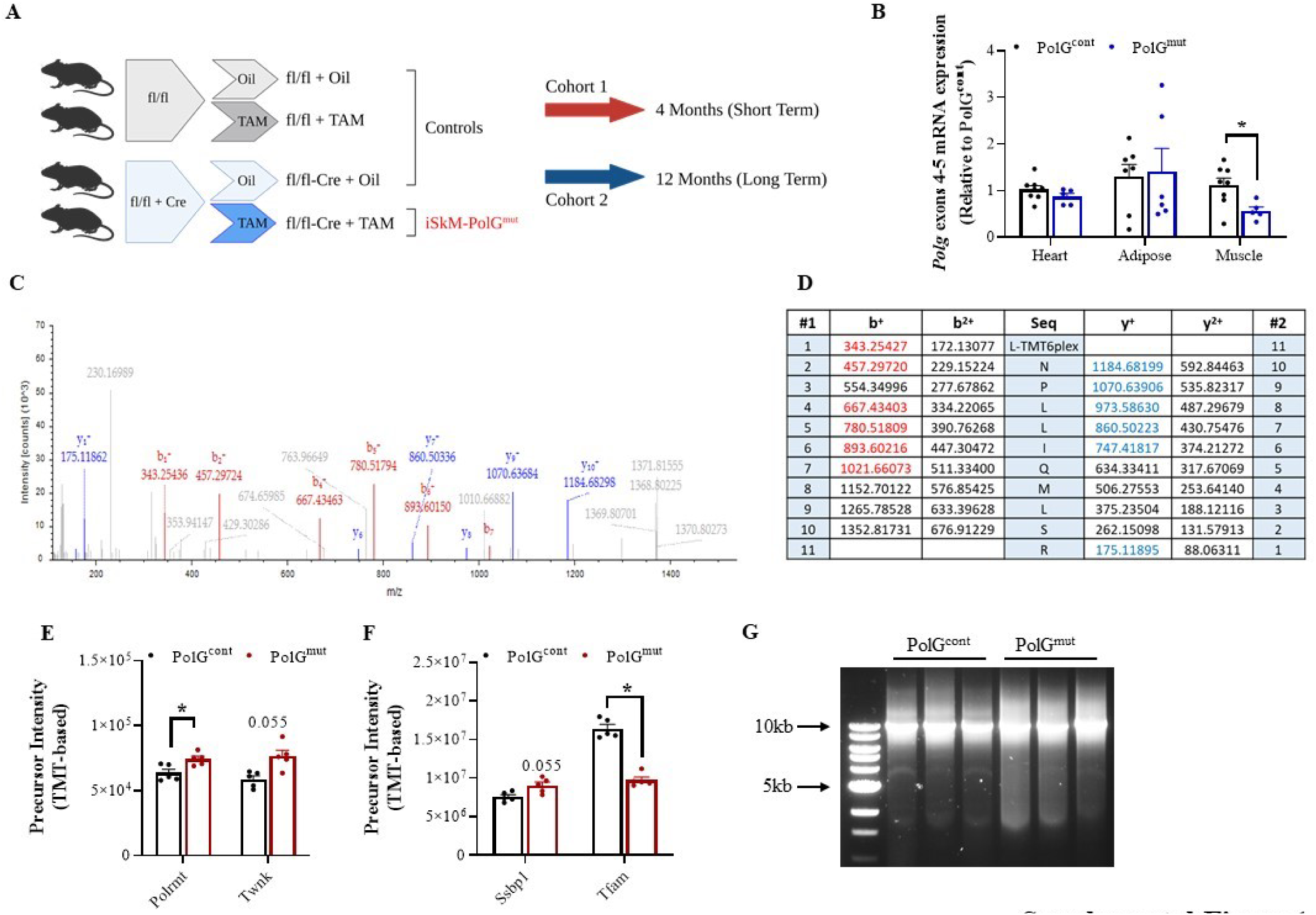
Generation of a Post-developmental Muscle Specific PolG Exonuclease Mutant. **(A)** Schematic outlining the mouse breeding strategy for (cohort 1 – 4 months, red; Cohort 2 – 12 months, blue), including controls (Tamoxifen control: fl/fl + TAM, Cre control: fl/fl + oil, Genetic control: fl/fl-Cre + oil) and muscle specific PolG mutant (PolG^mut^). **(B)** PolG gene expression in heart, adipose and TA muscle from PolG^cont^ and PolG^mut^ mice at Exon 4-5 (within floxed region) as determined by qPCR. **(C)** Spectral output of unique razor peptide from PolG, showing b- and y-ion series for specific m/z for identified peptide: **(D)** NPLLIQMLSR as used in proteomic analyses to quantify PolG abundance. Protein abundance in mitochondria isolated from PolG^cont^ and PolG^mut^ mice TA muscle, as determined by mass spectrometry proteomic analyses, of **(E)** Polrmt and Twnk, and **(F)** Ssbp1 and Tfam. **(G)** PCR amplified 10kb section (Taq634 site) from equivalent amounts of mtDNA isolated from PolG^cont^ and PolG^mut^ muscle. TA = Tibialis anterior Data are presented as mean ± SEM, n=5-10, * denotes p-value <0.05 between control and mutant samples as determined by unpaired, two-tailed Mann-Whitney t-tests.

**Supplemental Figure 2:**
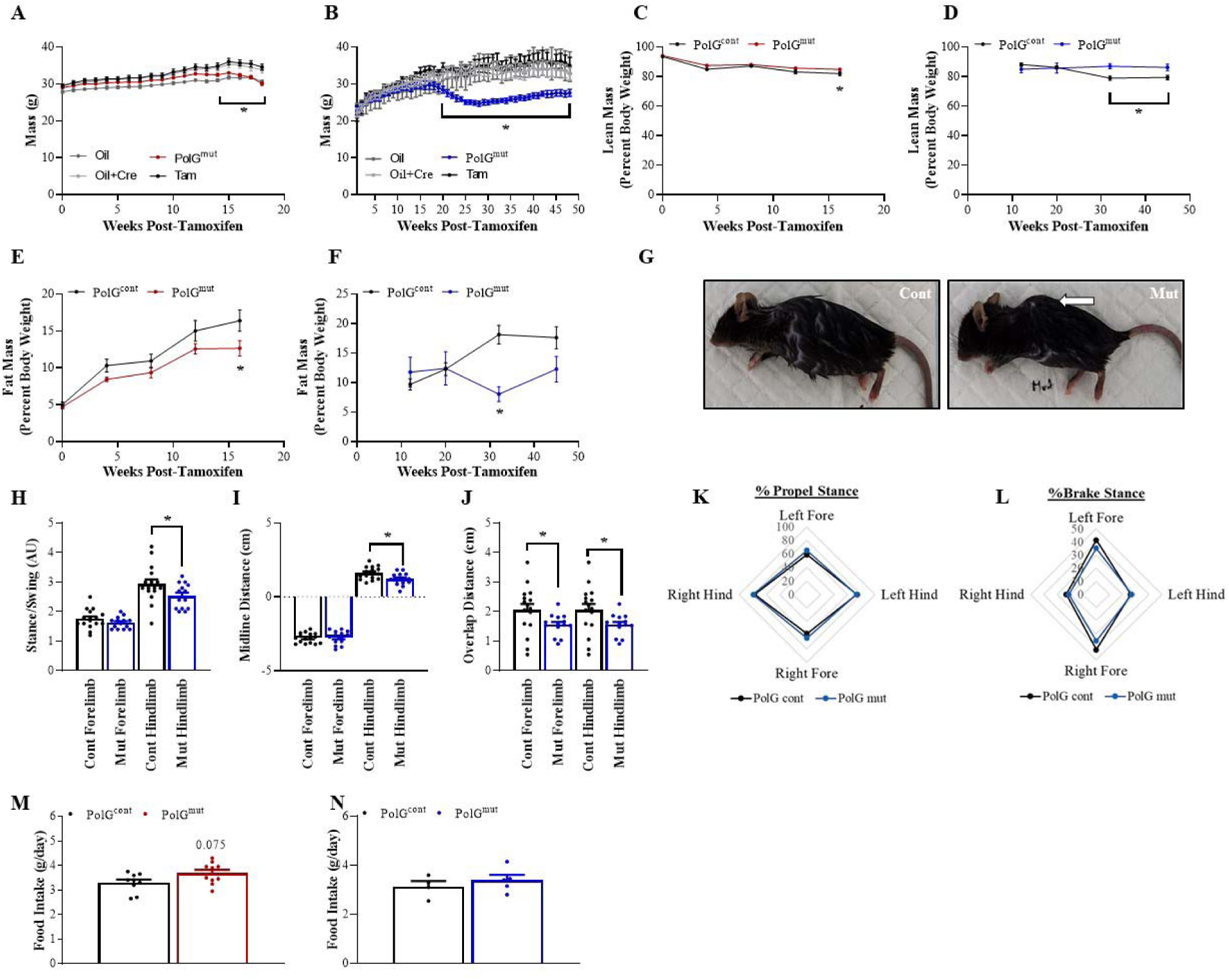
TNanoLC and Mass Spectrometry:he Effects of Muscle Specific loss of PolG Exonuclease Activity on Muscle Function and Whole Body Parameters. Cohorts of PolG^cont^ and PolG^mut^ mice were investigated for their metabolic health over both short (4 months, red lines/bars) and long (12 months, blue lines/bars) term time frames subsequent to the administration of tamoxifen. Progressive weekly weight (mass) measurements post-tamoxifen in **(A)** short-term PolG Controls and PolG^mut^ mice and **(B)** long term PolG^cont^ and PolG^mut^ mice. Progressive lean and fat mass as a percentage of total body weight in **(C&E)** short-term and **(D&F)** long term PolG^cont^ and PolG^mut^ mice. **(G)** Representative images indicating (arrow) the development of kyphosis in long term PolG^mut^ mice compared with PolG^cont^ mice. DigiGait analyses in long term PolG^cont^ and PolG^mut^ mice measuring **(H)** Stance/Swing ratio, **(I)** Midline Distance, **(J)** Overlap Distance, **(K)** %propel Stance, and **(L)** %Brake Stance. Daily food intake (over 24 hours) in **(M)** short-term and **(N)** long term PolG^cont^ and PolG^mut^ mice. Data are presented as mean ± SEM, n=5-10, * denotes p-value <0.05 between control and mutant samples as determined by repeated measures 2-way ANOVA with correction for multiple comparisons (A-F) or unpaired, two-tailed Mann-Whitney t-tests (H-N).

**Supplemental Figure 3:**
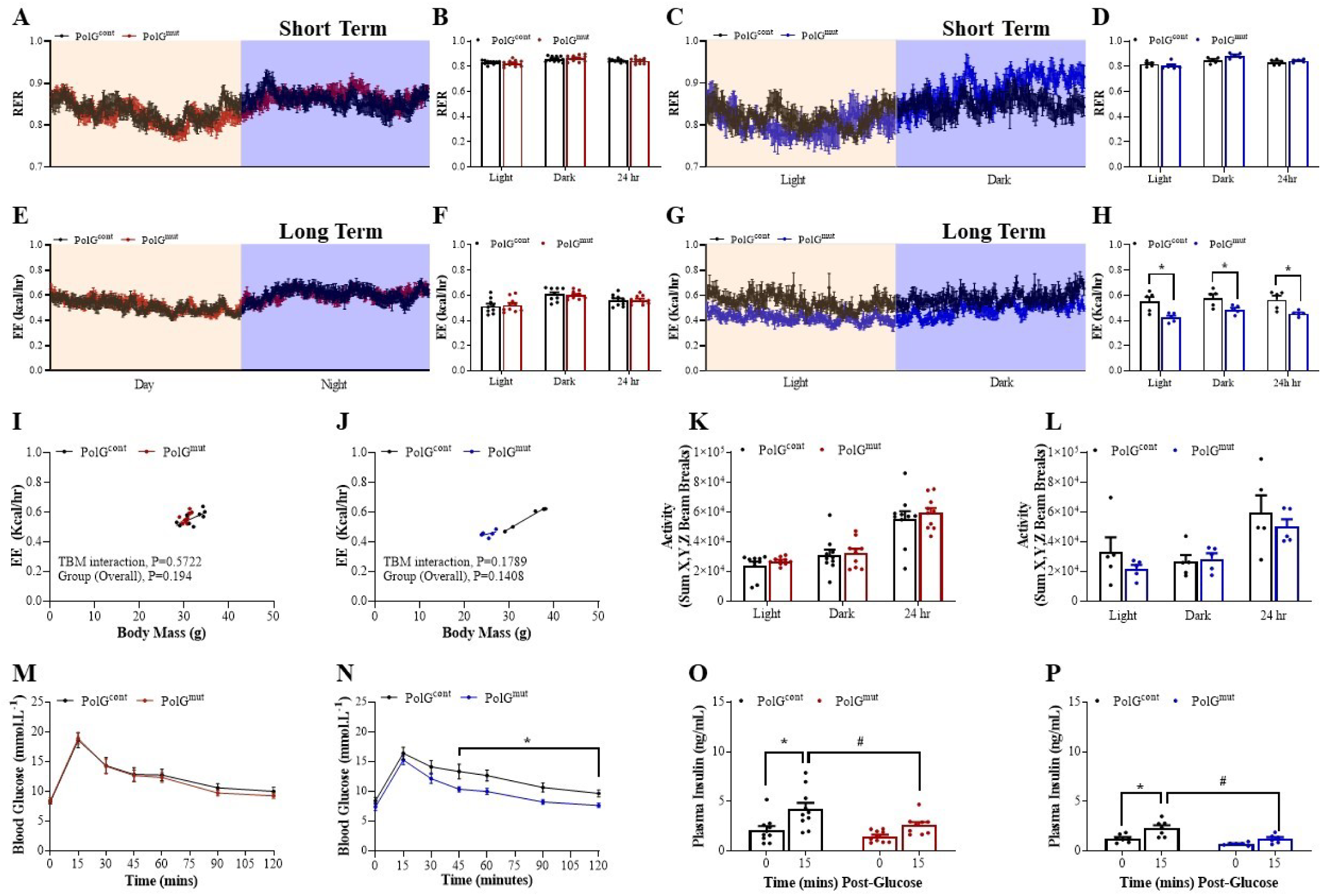
Whole Body Energy and Glucose Homeostasis in Muscle Specific PolG Exonuclease Mutant Mice. Cohorts of PolG^cont^ and PolG^mut^ mice were investigated for their metabolic health over both short (4 months, red lines/bars) and long (12 months, blue lines/bars) term time frames subsequent to the administration of tamoxifen. Mouse whole body energetics were analysed using the Promethion system, including RQ in **(A&B)** short-term and **(C&D)** long term cohorts, and EE in **(E&F)** short-term and **(G&H)** long term cohorts. ANCOVA analysis of energy expenditure (EE) as a function of body weight in the **(I)** short-term cohorts, and **(J)** in the long-term cohorts. Cumulative ambulatory movement (Sum X, Y, Z beam breaks) over a 24 hour period in **(K)** short-term cohorts and **(L)** long-term cohorts. Glucose tolerance tests were performed on both short and long-term cohorts, with subsequent data presented as line graph depicting change in plasma glucose as measured every 15 minutes over the 2 hours post-glucose gavage in **(M)** short-term and **(N)** long-term cohorts, with corresponding plasma insulin at 0 and 15 minute time points for **(O)** short-term and **(P)** long-term cohorts. RQ = respiratory quotient, EE = energy expenditure. Data are presented as mean ± SEM, n=5-10, * denotes p-value <0.05 between 0 and 15 minute points in insulin assay, # denotes p-value <0.05 between control and mutant samples and as determined by unpaired, two-tailed Mann-Whitney t-tests, repeated measures two-way ANOVA with correction for multiple comparisons, or ANCOVA.

**Supplemental Figure 4:**
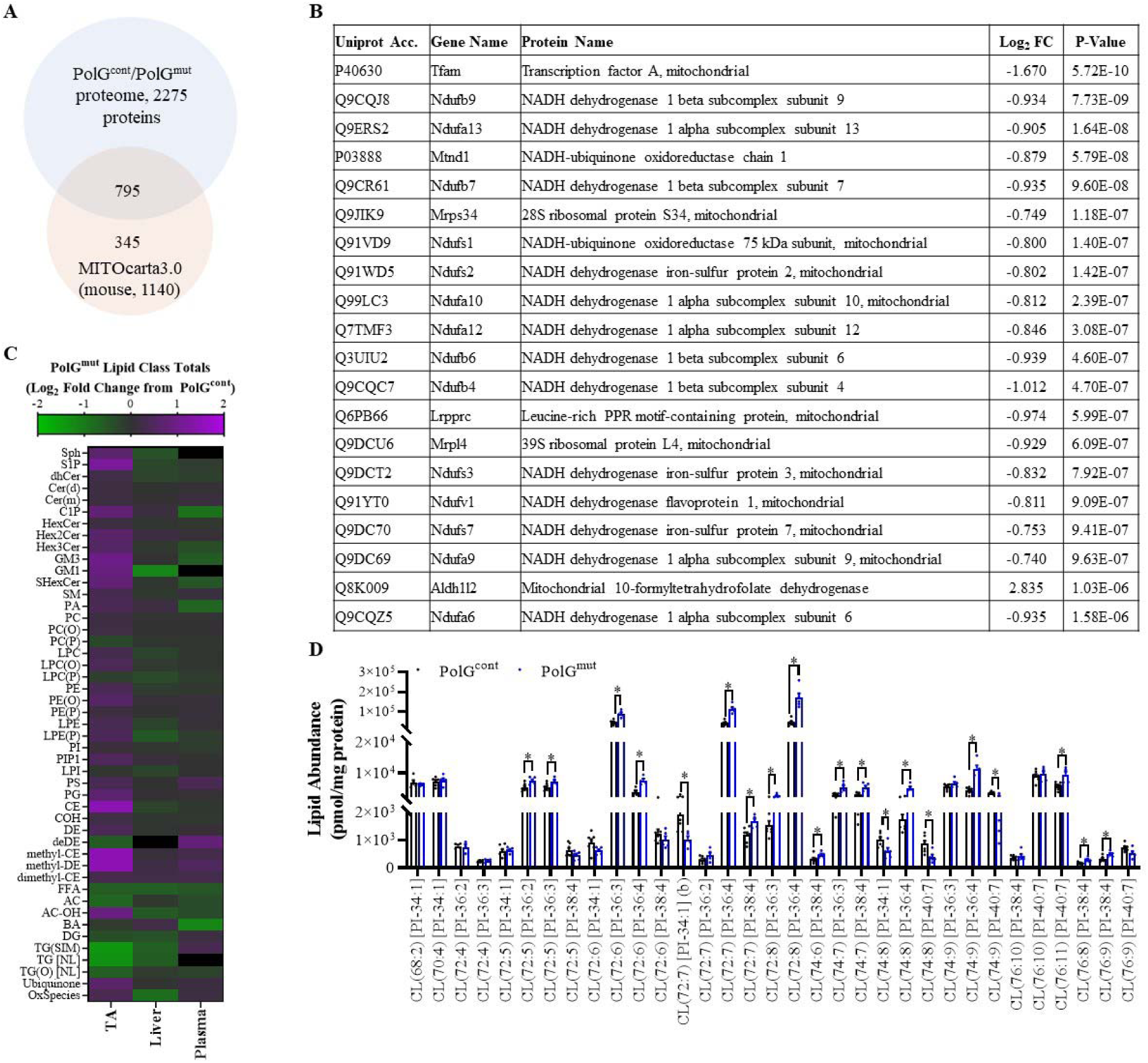
Mitochondrial Proteomic and Lipidomic Changes Induced by Muscle Specific loss of PolG Exonuclease Activity. Mitochondria were isolated from skeletal muscle of PolG^cont^ and PolG^mut^ mice that had completed the short-term experiments (4 months). Unbiased assessment of protein abundance in mitochondria was analysed using proteomics. **(A)** Proteomic analysis detected 1319 proteins in the PolG^cont^ mitochondria, and 1353 proteins in PolG^mut^ mitochondria, covering ∼70% of the known mitochondrial proteome as described in MITOcarta 3.0. **(B)** Table of the top 20 most significant differentially regulated proteins in PolG^mut^ compared with PolG^cont^ mice. **(C)** PolG^mut^ lipid class fold change from PolG^cont^ in TA muscle, liver and plasma (purple = increased, green = decreased in PolG^mut^). **(D)** Abundance of individual CL species (pmol/mg protein) in TA from PolG^cont^ and PolG^mut^ long term mice, where PI indicates Product Ion of the two Phosphatidic Acids with sum details of carbons and double bonds e.g. 72:8 = 2 x 36:4 (two acyl chains of 18:2). For proteomics n=6/group, and analysis of data were corrected using Benjamini-Hochberg FDR. Functional mitochondria data are presented as mean ± SEM, n=5-10, * denotes p-value <0.05 between control and mutant samples and as determined by unpaired, two-tailed Mann-Whitney t-tests.

**Supplemental Figure 5:**
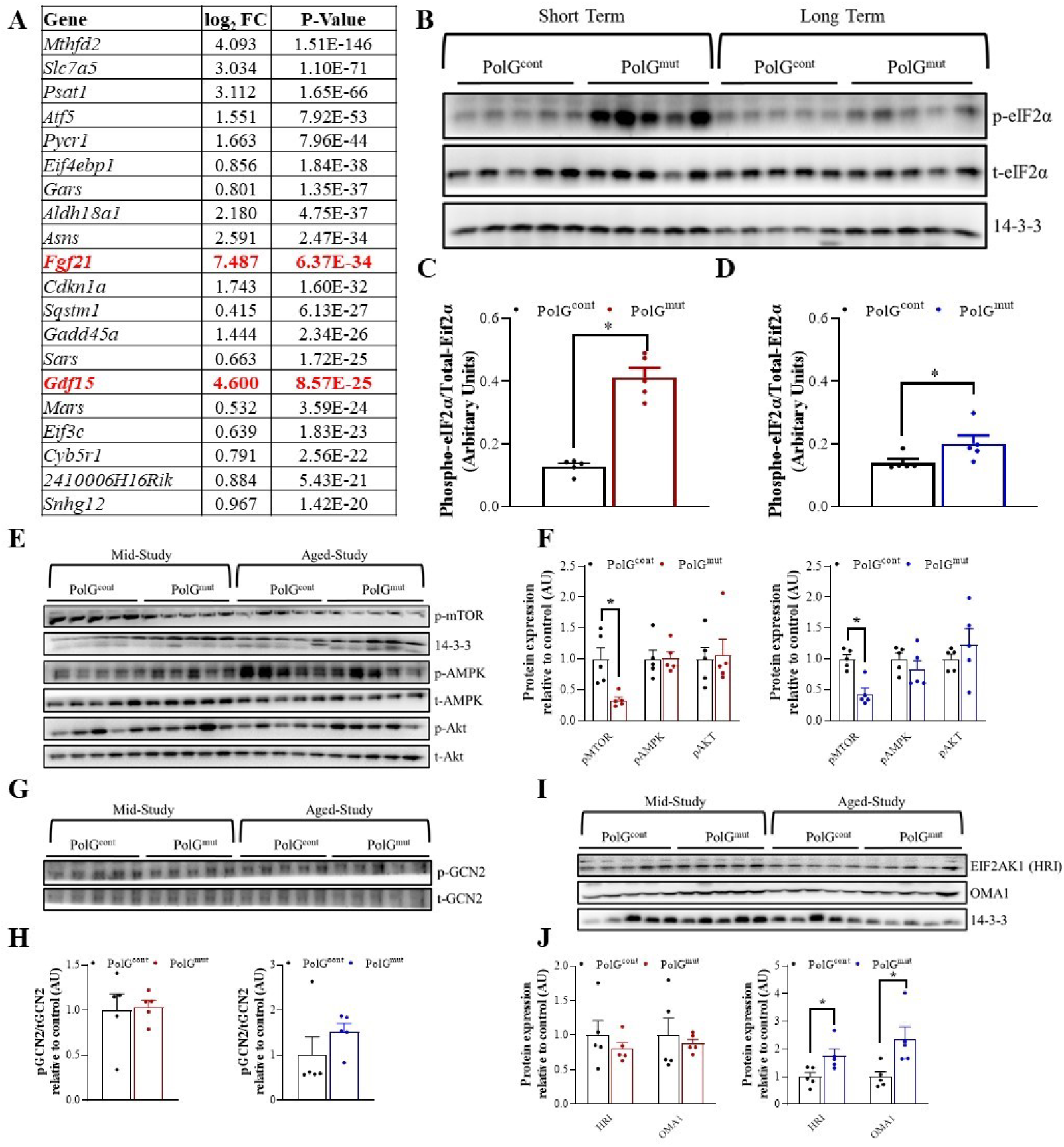
Transcriptional and Molecular Responses to Muscle Specific loss of PolG Exonuclease Activity. Unbiased assessment of RNA abundance was analysed from skeletal muscle of PolG^cont^ and PolG^mut^ short-term mice (4 months) using bulk RNA-sequencing. **(A)** The top 20 most significant differentially regulated genes (*Fgf21* and *Gdf15* highlighted in red). **(B)** Immunoblot for phospho/total eIF2α in muscle from PolG^cont^ and PolG^mut^ short-term (mid-study) and long term (aged-study) cohorts, and normalised to total eIF2α for **(C)** short-term cohort and **(D)** long term cohort. **(E-F)** Immunoblot for phospho mTOR and phospho/total Akt, AMPK in muscle from short and long-term cohorts normalised to either 14-3-3 (pMTOR) or total protein (AKT, AMPK). **(G-H)** Phospho/total GCN2 in muscle from PolG^cont^ and PolG^mut^ short and long-term cohorts and **(I-J)** HRI (EIF2AK1), OMA1 immunoblots and normalised to 14-3-3. For RNA-seq n=6/group, and analysis of data were corrected using Benjamini-Hochberg FDR. Immunoblot data are presented as mean ± SEM, n=5, * denotes p-value <0.05 between control and mutant samples as determined by unpaired, two-tailed Mann-Whitney t-tests.

**Supplemental Figure 6:**
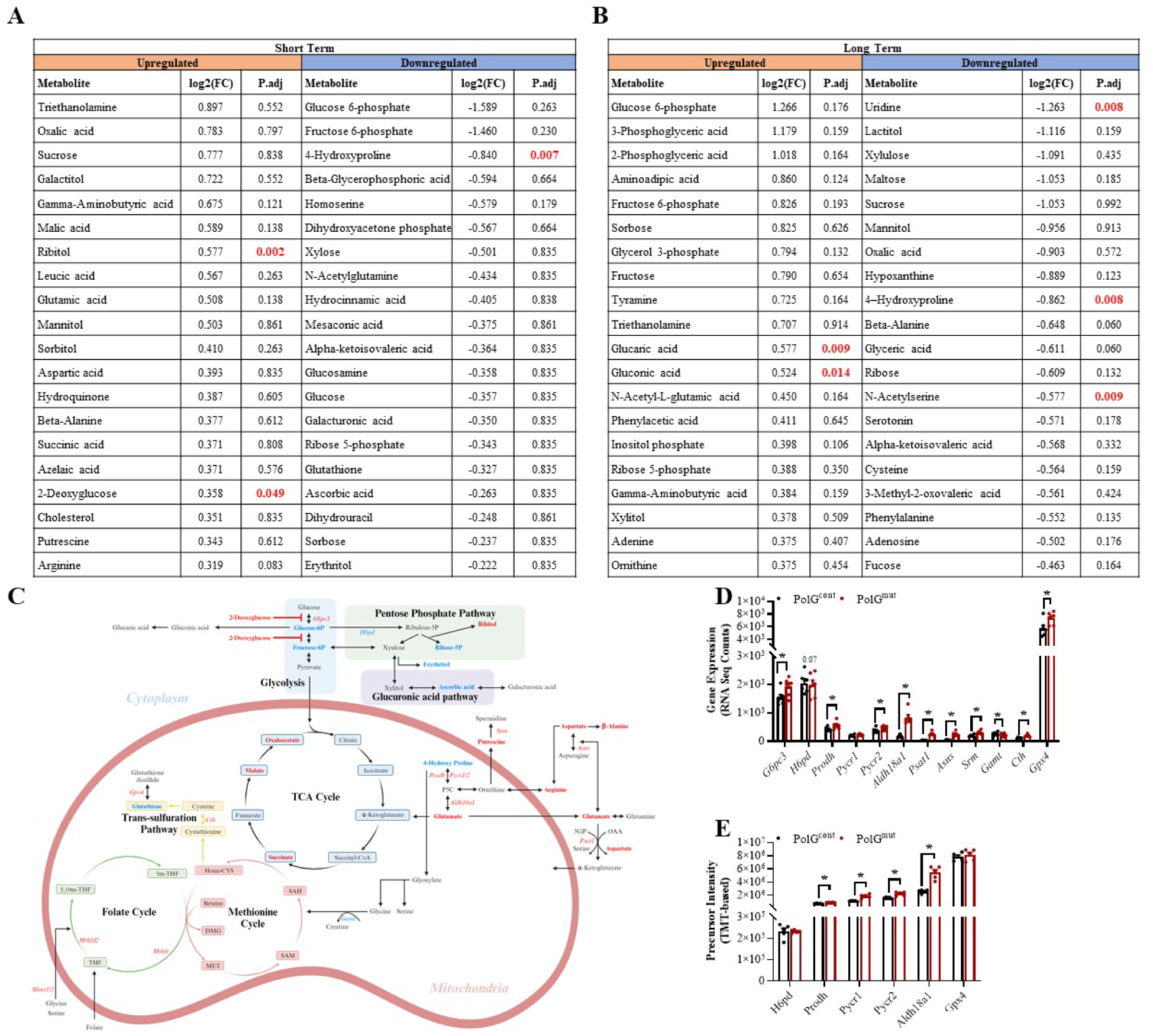
Metabolite adaptions in Muscle Specific PolG^mut^ Mice Mediated by Impaired Mitochondrial Function and the mtISR. Unbiased assessment of metabolite abundance was analysed from skeletal muscle of PolG^cont^ and PolG^mut^ short-term mice (4 months) and long-term mice. Table lists top 20 metabolites listed in order of largest fold-change (upregulated-beige, downregulated-blue) in PolG^mut^ mice compared with PolG^cont^ mice for **(A)** short-term and **(B)** long-term cohorts. Metabalomics data was normalised by median and log transformed using MetaboAnalyst 5.0 and analysis of data were corrected using Benjamini-Hochberg FDR, n=8/group (except PolG^mut^ long-term cohort, n=7). **(C)** Schematic of mammalian cytosolic/mitochondrial metabolic pathways, where red text denotes upregulated and blue text denotes downregulated genes (italic) and metabolites in short-term cohort. **(D)** Relevant metabolic genes detected by RNA-sequencing and **(E)** proteins detected by mitochondrial proteomics. Metabolomics data was normalised by median and log transformed using MetaboAnalyst 5.0, n=8/group (except PolG^mut^ long-term cohort, n=7). Analysis of data were corrected using Benjamini-Hochberg FDR. * denotes p-value <0.05 between control and mutant samples.

